# TGF-β-driven downregulation of the Wnt/β-Catenin transcription factor TCF7L2/TCF4 in PDGFRα^+^ fibroblasts

**DOI:** 10.1101/2020.01.05.895334

**Authors:** Osvaldo Contreras, Hesham Soliman, Marine Theret, Fabio M.V. Rossi, Enrique Brandan

## Abstract

Mesenchymal stromal/stem cells (MSCs) are multipotent progenitors essential ororganogenesis, tissue homeostasis, regeneration, and scar formation. Tissue injury upregulates TGF-β signaling, which modulates myofibroblast fate, extracellular matrix remodeling, and fibrosis. However, the molecular determinants of MSCs differentiation and survival remain poorly understood. The canonical Wnt Tcf/Lef transcription factors regulate development and stemness, but the mechanisms by which injury-induced cues modulate their expression remain underexplored. Here, we studied the cell-specific gene expression of Tcf/Lef and, more specifically, we investigated whether damage-induced TGF-β impairs the expression and function of TCF7L2, using several models of MSCs, including skeletal muscle fibro-adipogenic progenitors. We show that Tcf/Lefs are differentially expressed and that TGF-β reduces the expression of TCF7L2 in MSCs but not in myoblasts. We also found that the ubiquitin-proteasome system regulates TCF7L2 proteostasis and participates in TGF-β-mediated TCF7L2 protein downregulation. Finally, we show that TGF-β requires HDACs activity to repress the expression of TCF7L2. Thus, our work found a novel interplay between TGF-β and Wnt canonical signaling cascades in PDGFRα^+^ fibroblasts and suggests that this mechanism could be targeted in tissue repa ir and regeneration.

**Summary statement:** TGF-β signaling suppresses the expression of the Wnt transcription factor TCF7L2 and compromises TCF7L2-dependent functions in tissue-resident PDGFRα^+^ fibroblasts.

## INTRODUCTION

The remarkable long-term capacity of adult skeletal muscle to grow and regenerate is largely based on the presence of tissue-resident muscle stem cells (MuSCs) (formerly called satellite cells) (Lepper et al., 2011; Murphy et al., 2011; Sambasivan et al., 2011). However, another tissue-resident population of quiescent mesenchymal stromal/stem cells (MSCs) is also essential for effective regeneration and maintenance of the skeletal muscle and its connective tissue (CT). Following acute damage, PDGFRα^+^ MSCs provide regenerative cues, regulate extracellular matrix (ECM) remodeling, and MuSCs behavior, thus facilitating muscle regeneration and maintenance (Contreras, 2019d; Heredia et al., 2013; Joe et al., 2010; Scott et al., 2019; Uezumi et al., 2010; Wosczyna et al., 2019). These stromal mesenchymal progenitors are also called fibro-adipogenic progenitors (FAPs) in mouse and human skeletal muscle (Agley et al., 2013; Dulauroy et al., 2012; Uezumi et al., 2014a; Vallecillo-García et al., 2017; Wosczyna and Rando, 2018). Muscle MSCs have the potential to differentiate *in vivo* and *in vitro* into myofibroblasts, adipocytes, chondrogenic, and osteogenic cells (Agley et al., 2013; Contreras et al., 2019c; Oishi et al., 2013; Uezumi et al., 2014b; Uezumi et al., 2010; 2011; Wosczyna et al., 2012). PDGFRα^+^ mesenchymal progenitors are found in most mammalian tissues, where they regulate tissue homeostasis, repair, and regeneration (Carr et al., 2019; Riquelme-Guzmán and Contreras, 2020; Lemos and Duffield, 2018; Lynch and Watt, 2018; Rognoni et al., 2018). Despite their required normal activity during muscle regeneration, we and others have suggested dysregulated behavior of these mesenchymal precursor cells in models of chronic muscle damage, muscular dystrophy (MD), neurodegenerative diseases, and aging (Acuña et al., 2014; Contreras et al., 2016; Contreras et al., 2019c; González et al., 2017; Ieronimakis et al., 2016; Kopinke et al., 2017; Madaro et al., 2018; Mahmoudi et al., 2019; Lemos et al., 2015; Lukjanenko et al., 2019; Uezumi et al., 2014a). Fibrosing disorders are a common outcome of the dysregulation of these fibroblastic cells, which include non-malignant fibroproliferative diseases with high morbidity and mortality (Lemos and Duffield, 2018; Wynn and Ramalingam, 2012). However, the participation of mesenchymal progenitors in repair and regeneration remains poorly understood.

The Wnt signaling pathway has a key role in many aspects of developmental biology, tissue homeostasis, stem cells fate, organogenesis, cancer, and tissue fibrogenesis. Secreted Wnt ligands, which are 19 secreted lipid-modified glycoproteins, bind to the Frizzled receptors and its co-receptor LRP5/6 on the cell surface to initiate a signaling pathway that regulates the proteostasis of cytoplasmic β-catenin. In the absence of canonical Wnt ligands, β-catenin is a target for the UPS-mediated degradation (Aberle et al., 1997). Once the canonical cascade starts, the accumulation of β-catenin leads to its translocation to the nucleus, where it recognizes and binds the T-cell factor (TCF) or Lymphoid enhancer (LEF) transcription factors (TFs), but it also recruits transcriptional partners and chromatin remodeling complexes which in concert regulate the expression of Tcf/Lef target genes. These TFs recognize Tcf/Lef-binding elements on its target genes to regulate the expression of thousands of genes (Cadigan and Waterman, 2012; Clevers, 2006; Schuijers et al., 2014). Therefore, the Tcf/Lef TFs are the final effectors of the canonical Wnt/β-catenin signaling cascade in metazoans (Nusse and Clevers, 2017; Schuijers et al., 2014; Tang et al., 2008). Mammalian cells express four Tcf/Lef protein-coding genes: Tcf1 (Tcf7), Lef1, Tcf7l1 (formerly named Tcf3), and Tcf7l2 (formerly named Tcf4) (Cadigan and Waterman, 2012; van de Wetering et al., 1991). They have critical roles regulating the body plan establishment, cell fate specification, proliferation, survival, and differentiation, which are predominant features in fast and constantly renewing tissues (Clevers, 2006; Korinek et al., 1998). Accumulating evidence indicates that pathologically activated canonical Wnt signaling plays a major role in the pathogenesis of fibrosis in multiple tissues (Chilosi et al., 2003; Cisternas et al., 2014; Colwell et al., 2006; Cosin-Roger et al., 2019; He et al., 2009; He et al., 2010; Henderson et al., 2010; Konigsho et al., 2008; Liu et al., 2009; Surendran et al., 2002; Trensz et al., 2010; Wei et al., 2011). For example, the conditional genetic loss of β-catenin in cardiac fibroblast-myofibroblast lineages (Tcf21^+^ or Periostin^+^) causes the reduction of interstitial fibrosis and attenuates heart hypertrophy induced by cardiac pressure overload (Xiang et al., 2017). Furthermore, upregulated Wnt activity during aging accounts in part for the declining regenerative potential of muscle with age and increased fibrosis (Brack et al., 2007). Transgenic mice overexpressing canonical Wnt-10b demonstrated that activation of this pathway is sufficient to induce fibrosis *in vivo* (Akhmetshina et al., 2012). Mechanistically, it has been demonstrated that Wnt/β-catenin pathway regulates the expression of several ECM genes in fibroblasts from different organs (Akhmetshina et al., 2012; Hamburg-Shields et al., 2015; Xiang et al., 2017). Altogether, these studies suggest that Wnt/β-catenin signaling in stromal cells is required for pathogenic ECM gene expression and collagen deposition during fibrogenesis.

Transforming growth factor type-beta (TGF-β) regulates the differentiation program of a variety of cell types (for review see David and Massagué, 2018; Derynck and Budi, 2019), including MSCs. Mechanistically, TGF-β signaling stimulates PDGFRα^+^ cells proliferation, ECM production, and myofibroblast differentiation (Contreras et al., 2019b; Lemos et al., 2015; Uezumi et al., 2011). Three TGF-β isoforms -TGF-β1, TGF-β2, and TGF-β3-are expressed in mammals (Massagué, 1998). These isoforms and their mediated signaling pathways are exacerbated in Duchenne muscular dystrophy (DMD) patients (Bernasconi et al., 1999; Smith and Barton, 2018), skeletal muscle of the *mdx* mice (Gosselin et al., 2004; Lemos et al., 2015), and during fibrogenesis in several organs (Kim et al., 2018). Thus, it is recognized that dysregulated TGF-β activity is a driver for reduced muscle regeneration, impaired tissue function and fibrosis (Juban et al., 2018; Mann et al., 2011; Pessina et al., 2015; Vidal et al., 2008). Interestingly, the inhibition of TGF-β improves the pathophysiology of MD (Accornero et al., 2014; Acuña et al., 2014; Ceco and McNally, 2013; Cohn et al., 2007; Danna et al., 2014).

Here, we show that Tcf/Lef members are differentially expressed in tissue-resident PDGFRα^+^ cells and demonstrate that *Tcf7l2* is a novel TGF-β target gene. The ubiquitin-proteasome system (UPS) participates, both, in modulating the TCF7L2 proteostasis and TGF-β-mediated downregulation of TCF7L2. We also found that TGF-β requires the activity of HDACs to repress *Tcf7l2* gene expression and to alter Wnt signaling gene signature. Finally, these effects were not observed in myogenic cells, suggesting that this crosstalk is fibroblast/MSC specific.

## RESULTS

### Wnt canonical Tcf/Lef transcription factors are differentially expressed in PDGFRα^+^ mesenchymal progenitor cells and fibroblasts

We first determined the relative expression of the Wnt T-cell factor/Lef members in the mesenchymal stromal/stem cell line C3H/10T1/2 and mouse embryonic fibroblasts (MEFs) (Fig. 1A). *Tcf7l1* and *Tcf7l2* were the two most highly expressed members of this family, while *Lef1* and *Tcf7* were almost not expressed in MSCs or MEFs (Fig. 1A). Then, we evaluated the expression of these transcription factors in *ex vivo* FACS-isolated skeletal muscle PDGFRα-H2BEGFP^+^ FAPs (Fig. 1B, Fig. S1A). Our results using muscle PDGFRα^+^ cells further corroborated the observations described above (Fig. 1C). We further corroborated these results using RNA-seq data of mononuclear muscle fractions (Scott et al., 2019). We focused on a Lin^−^(CD31^−^CD45^−^Ter119^−^) LY6A/Sca1^+^ population and observed that *Tcf7l2* and *Tcf7l1* transcripts were enriched in the Lin^-^Sca1^+^ fraction (Fig. 1D,E). Thus, these results suggest that among the Wnt Tcf/Lef members, *Tcf7l1* and *Tcf7l2* but not *Lef1* or *Tcf7*, are highly expressed in PDGFRα^+^ cells. We next evaluated TCF7L2 protein expression in skeletal muscle tissue and MSCs. The protein products detected by western blotting in most tissues and cells are ∼78 and ∼58 kDa in size (Tang et al., 2008; Weise et al., 2010). We found that these TCF7L2 protein isoforms are present in limb muscles and MSCs, representing the E and M/S isoforms (Fig. 1F, Fig. S1B) (Jin, 2016). We also found that the TCF7L2 TF is predominantly expressed in the nucleus of muscle-resident interstitial cells of the undamaged diaphragm muscle, isolated EGFP^+^ FAPs from PDGFRα^H2BEGFP^ mice, *mdx*;PDGFRα^H2BEGFP^ dystrophic mice, and MEFs (Fig. 1G, Fig. S1B-D). The tissue-specific expression of the Tcf/Lef members and their relative expression in MSCs was further corroborated by additional single-cell RNAseq data (Fig. S1F,G) (The Tabula Muris Consortium et al., 2018). Next, we evaluated the expression of TCF7L2 in transiently activated PDGFRα^+^ FAPs and confirmed that TCF7L2 TF is present in these cells three days after notexin injury (Fig. 1H,I). Stromal PDGFRα^+^ fibroblasts are also found in the heart, playing supportive roles in cardiac development and repair (Chong et al., 2011; Farbehi et al., 2019; Fu et al., 2018; Furtado et al., 2016; Soliman et al., 2020; Tallquist and Molkentin, 2017). Similar to what we found in MSCs cell lines and muscle PDGFRα^+^ FAPs, *Tcf7l1* and *Tcf7l2* are the two highest expressed members of this family in cardiac PDGFRα^+^ fibroblasts whereas *Lef1* and *Tcf7* have very low expression levels (Fig. 1J). We further corroborated these results using RNA-seq data from cardiac fibroblasts (Fu et al., 2018). We also found that *Tcf7l2* and *Tcf7l1* transcripts were enriched in the heart PDGFRα^+^-Tcf21^+^ fibroblast population (Fig. 1K). Similar to muscle, both E and M/S TCF7L2 protein isoforms were also found in cardiac tissue and PDGFRα^+^ fibroblasts (Fig. 1M, N). Altogether, these results establish that specific Tcf/Lef transcription factors, namely TCF7L2/TCF7L1, are highly expressed in PDGFRα^+^ MSCs and fibroblasts as well as in total skeletal muscle and cardiac tissue.

### Dynamics of TCF7L2 expression in stromal fibroblasts during regeneration and repair

Us and other investigators demonstrated that sub-populations of stromal MSCs co-express both TCF7L2 and PDGFRα during skeletal muscle development and also in adulthood (Contreras et al., 2016; Contreras et al., 2019a,b; Murphy et al., 2011; Vallecillo-García et al., 2017). As a consequence of the stromal expansion caused by injury, both TCF7L2 and PDGFRα protein levels are increased in dystrophic tissue, denervated muscle, and after chronic barium chloride-induced damage (Contreras et al., 2016). Interestingly, the number of TCF7L2^+^ cells and TCF7L2 bulk protein levels correlate with the extension of damage, fibrosis, and TGF-β levels in muscles of the *mdx* mice (Contreras et al., 2016; Contreras et al., 2019a). Additionally, the expansion of TCF7L2^+^ MSCs in human regenerating muscle closely associates with regenerating myofibers *in vivo* (Mackey et al., 2017). Thus, to study whether the expression of TCF7L2 changes during regeneration and repair, we used two previously described muscle injury models (Uezumi et al., 2010; Kopinke et al., 2017). First, we aimed to study whole tissue TCF7L2 protein levels during skeletal muscle regeneration following glycerol-induced acute damage, a model that leads to transient expansion of PDGFRα^+^ cells (Contreras et al., 2019c; Kopinke et al., 2017). As expected, glycerol acute damage caused an increase in total TCF7L2 protein levels, which transiently peak at day 3 following glycerol intramuscular injection (Fig. 2A,B). PDGFRα bulk protein levels increased with similar kinetics of expression as TCF7L2 after acute injury (Fig. 2A,B). We also found that β-Catenin, the transcriptional partner of TCF7L2, increased after glycerol damage (Fig. 2A,B). Myosin heavy chain expression, a marker of regenerating myofibers, peaked at day 7 after damage (Fig. 2A,B). As mentioned above, we previously described that TCF7L2^+^ stromal cells and TCF7L2 bulk expression increases in the dystrophic diaphragm (Fig. 2C,D) (Contreras et al., 2016). Intriguingly, we found a reduction in the expression levels of TCF7L2 in the expanded population of TCF7L2^+^ cells in the dystrophic diaphragm, where the proportion of TCF7L2^+^ cells expressing low levels of TCF7L2 was larger compared to undamaged muscle. In the latter case, TCF7L2 expression was generally high (Fig. 2C). To further explore our observation, we used confocal microscopy and z-stack reconstructions of transverse muscle sections to analyze the cellular amount of TCF7L2 in TCF7L2^+^ cells in the dystrophic diaphragm (Fig. 2C,E, Fig S1H). TCF7L2^high^ cells were distributed throughout the muscle interstitium, consistent with previous studies of fibroblast distribution (Fig. 2C, Fig. S1H) (Contreras et al., 2016; Contreras et al., 2019c; Mathew et al., 2011; Merrell et al., 2015; Murphy et al., 2011). On the other hand, TCF7L2^medium/low^-expressing cell number expanded, and their expansion corresponded to increased phosphorylation of Smad3, an index of activated TGF-β signaling in interstitial cells and muscle fibers, as well as in collagen type 1-enriched areas (Fig. 2F-H, Fig. S1H). We further corroborated our results using RNA-seq data from undamaged PDGFRα^+^ cardiac fibroblasts and after myocardial infarction (MI) (Fu et al., 2018). Proliferation, TGF-β, and Matrisome-ECM-modifying genes were most highly induced at the initial stages (days 3 and 7) but then they were downregulated 2 and 4 weeks after MI (Fig. S2). However, we observed that the expression of *Tcf7l2* and *Tcf7l1* was highest in quiescent fibroblasts and became downregulated upon fibroblast activation, while expression of *Lef1* followed the opposite trend (Fig. S2). *Pdgfra* and *Tcf21* followed a similar kinetic pattern to that of *Tcf7l2* and *Tcf7l1*. Altogether, these results suggest that the magnitude of TCF7L2^medium/low^ cells expansion depends on the extent of tissue inflammation, TGF-β levels, damage, and fibrosis, and hence they were abundant in the inflammatory dystrophic model and after MI.

**Fig. 1.**
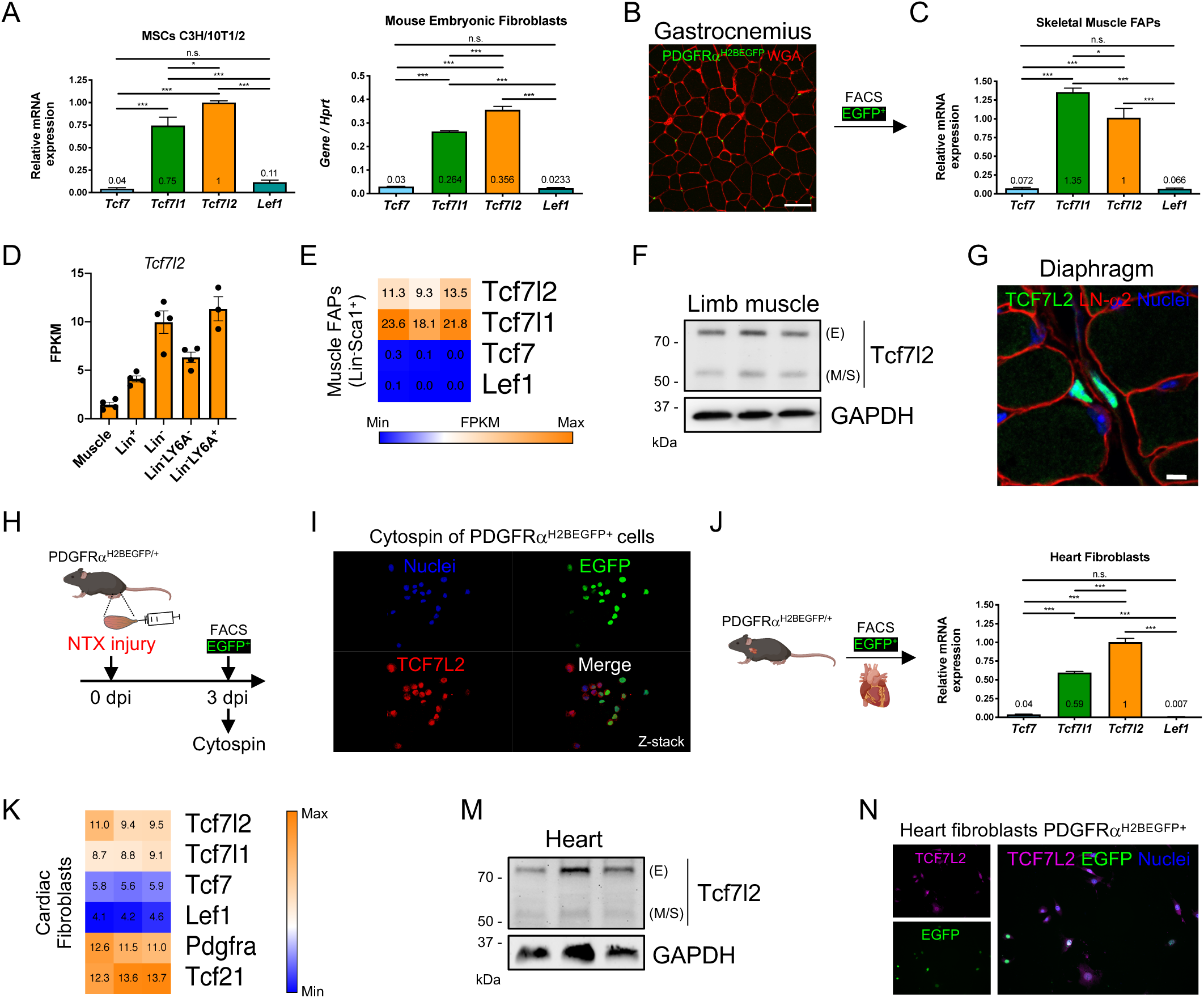
Differential expression of Tcf/Lef transcription factors in mesenchymal progenitor cells and fibroblasts. (A) *Tcf7, Tcf7l1, Tcf7l2*, and *Lef1* mRNA expression levels were analyzed by quantitative PCR in C3H/10T1/2 cells and MEFs in growing conditions. \*\*\**P*<0.001; \**P*<0.05; n.s., not significant by one-way ANOVA with Dunnett’s post-test; *n*=4. (B) Skeletal muscle FAPs were FACS-isolated from the PDGFRα^H2BEGFP^ mice. WGA staining labels lectins of the ECM. Scale bar: 50 μm. (C) *Tcf7, Tcf7l1, Tcf7l2*, and *Lef1* mRNA expression levels were analyzed by quantitative PCR in EGFP^+^ FAPs in growing conditions. \*\*\**P*<0.001; \**P*<0.05; n.s., not significant by one-way ANOVA with Dunnett’s post-test; *n*=3. (D) *Tcf7l2* transcript abundance in the different muscle cell fractions. (n = 3–4). (E) Heat map showing gene expression of Tcf/Lef in muscle Lin^-^Sca1^+^ population (FAPs). (F) Representative western blot analysis showing TCF7L2 protein levels in gastrocnemius muscle. GAPDH was used as the loading control. (G) Confocal image of TCF7L2 immunofluorescence. Laminin-α2 (LN-α2) (*red*) and nuclei (Hoechst in *blue*) were also stained. Scale bar: 10 μm. (H) Strategy used to isolate transit amplifying EGFP+ FAPs at day 3 post NTX TA injury. (I) Z-stack confocal image of cytospined EGFP+ FAPs showing TCF7L2 nuclear expression. (J) *Tcf7, Tcf7l1, Tcf7l2*, and *Lef1* mRNA expression levels were analyzed by quantitative PCR in cardiac fibroblasts in growing conditions. \*\*\**P*<0.001; n.s., not significant by one-way ANOVA with Dunnett’s post-test; *n*=3. (K) Heat map showing gene expression levels of Tcf/Lef in cardiac fibroblasts. Known cardiac fibroblast marker genes (*Pdgfra* and *Tcf21*) are also shown. Each column represents an individual uninjured cardiac fibroblast (*n*=3). (M) Representative western blot analysis showing TCF7L2 protein levels in whole cardiac tissue. GAPDH was used as the loading control. (N) Immunofluorescence of TCF7L2 (*magenta*) in FACS-isolated heart PDGFRα-EGFP+ cells in growing conditions.

**Fig. 2.**
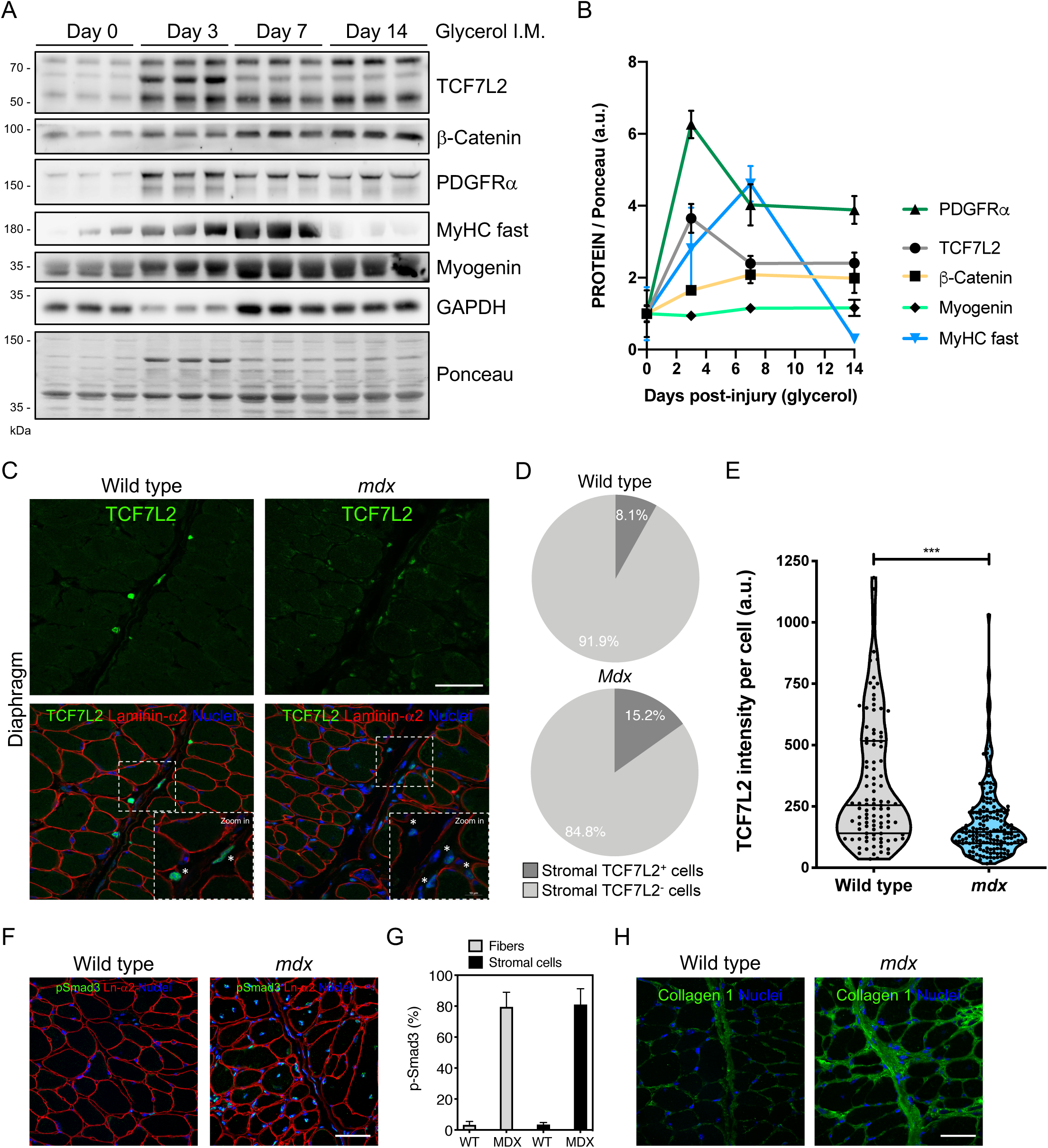
Dynamics of TCF7L2 expression in the stromal compartment during skeletal muscle regeneration and repair. (A) Representative western blot analysis showing TCF7L2, β-Catenin, PDGFRα, MyHC fast, Myogenin, and GAPDH in TA muscle after damage with glycerol at different time points (0, 3, 7 and 14 day). Ponceau red was used as the loading control. (B) Quantification of TCF7L2, β-Catenin, PDGFRα, MyHC fast, and Myogenin protein expression. *n*=3. (C) Z-stack confocal images showing the localization of TCF7L2^+^ cells in diaphragm muscle sections of adult wild-type and from the *mdx* mice. Scale bars: 50 μm and 10 μm. (D) Pie charts showing the increase of TCF7L2^+^ cells in dystrophic diaphragm. (E) Quantification of fluorescence intensity of TCF7L2 in stromal TCF7L2^+^ cells. ***P<0.001; by two-tailed Student’s t-test. n=4. (F) Representative confocal image showing the localization of phosphorylated-Smad3^+^ cells in diaphragm muscle sections of adult wild-type and *mdx* mice. Scale bar: 50 μm. (G) Quantification of the percentage of phospho-Smad3^+^ cells. n=4. (H) Representative confocal image showing the increase of ECM collagen type 1 immunostaining in diaphragm of *mdx* compared to wild-type mice (*n*=3).

### TGF-β signaling downregulates the expression of the WNT TCF7L2 transcription factor in PDGFRα^+^ FAPs, MSCs and fibroblasts

Chronic upregulation of TGF-β is found in several models of organ damage, where it is known to regulate the severity of tissue fibrosis. Activated extracellular TGF-β not only promotes MSCs survival and proliferation but also primes these progenitor cells to become myofibroblasts (Contreras et al., 2019b, 2019c; Cho et al., 2018; Kim et al., 2018; Lemos et al., 2015). TCF7L2^+^ cells expand near CD68^+^ macrophages in dystrophic muscles (one of the major cell sources of TGF-β in DMD) (Contreras et al., 2016; Juban et al., 2018; Tidball and Villalta, 2010). Our previous results suggest that the cell-specific expression of TCF7L2 negatively correlates with damage and inflammation in dystrophic muscle. Hence, damage-associated signaling pathways might regulate TCF7L2 expression and function endogenously during tissue repair. Therefore, we aimed to investigate the role of TGF-β signaling on the expression of TCF7L2 in fibroblast-related cell types and tissue-resident PDGFRα^+^ cells. We first used the multipotent mesenchymal progenitor cell line C3H/10T1/2 as an *in vitro* model of MSCs, because these cells have been extensively used to study mesenchymal biology (Braun et al., 1989; Contreras et al., 2019c; Reznikoff et al., 1973; Riquelme-Guzmán, et al., 2018; Singh et al., 2003). TGF-β1 diminished TCF7L2 protein expression in C3H/10T1/2 MSCs in a concentration-dependent manner, with effects already noticeable at the pathophysiological concentration of 0.5 ng/ml (Fig. 3A,B, Fig. S3A,B). We confirmed that C3H/10T1/2 cells respond to TGF-β1 in a concentration dependent-manner, by increasing the expression of ECM-related proteins (fibronectin, β1-Integrin, CCN2/CTGF) and αSMA (myofibroblasts marker), as well as by reducing the expression of PDGFRα (Fig. 3A,B) (Contreras et al., 2019c). Moreover, TGF-β1-mediated reduction of TCF7L2 expression began 8 h after stimulation, reaching a maximum at 24 and 48 h. At this point the expression of TCF7L2 had decreased by ∼85 per cent (Fig. 3C,D). Similar effects were also seen in the embryonic fibroblast NIH-3T3 cell line (Fig. 3E,F, Fig. S3A,B). Taken together, these results suggest that TGF-β downregulates TCF7L2 protein expression in a concentration- and time-dependent manner in MSCs and fibroblasts. Consistently with the above-mentioned results, we found that TGF-β1 reduced the expression of TCF7L2 by ∼50% in FACS-isolated muscle PDGFRα^+^ FAPs (Fig. 3G,H). As expected, TGF-β increased the expression of the ECM protein fibronectin (Fig. 3G) (Uezumi et al., 2010; Contreras et al., 2019c). Taken together, our results indicate that, concurrently with the induction of fibroblast activation and differentiation, TGF-β inhibits the expression of TCF7L2 in skeletal muscle PDGFRα^+^ FAPs, and in two different mesenchymal cell lines, C3H/10T1/2 and NIH-3T3 fibroblasts.

**Fig. 3.**
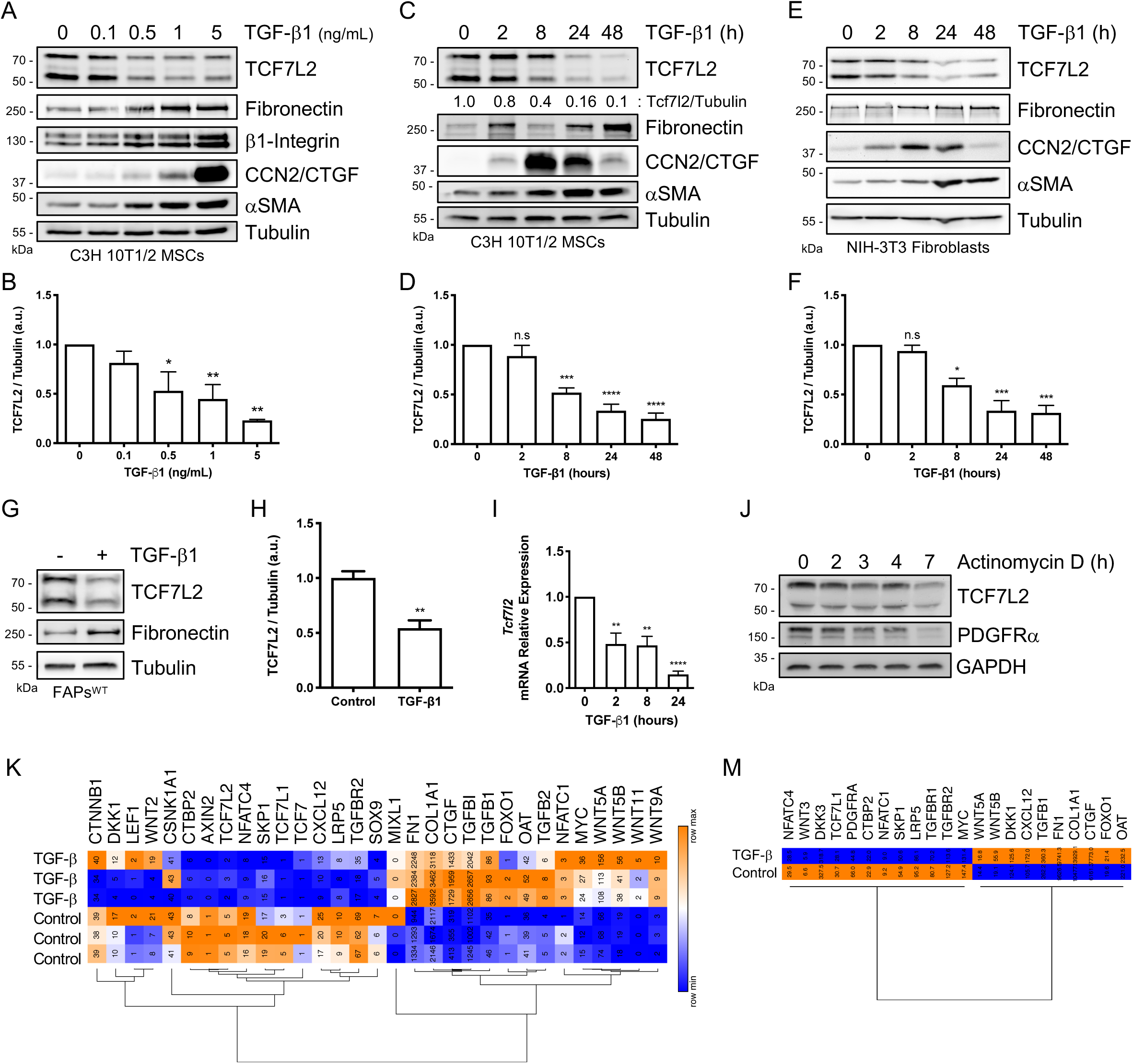
TGF-β signaling downregulates the expression of the Wnt TF TCF7L2 in mesenchymal progenitors and fibroblasts. (A) Representative western blot analysis showing TCF7L2, fibronectin, β1-Integrin, CCN2/CTGF, and αSMA/ACTA2 expression levels in C3H/10T1/2 MSCs after treatment with different concentrations of TGF-β1 for 24 h. Tubulin was used as the loading control. (B) Quantification of TCF7L2 protein expression. \*\**P*<0.005; \**P*<0.05 by one-way ANOVA with Dunnett’s post-test; *n*=4. (C,E) Representative western blot analysis showing TCF7L2, fibronectin, CCN2/CTGF, and αSMA/ACTA2 expression levels in (C) C3H/10T1/2 MSCs and (E) NIH-3T3 fibroblasts after treatment with TGF-β1 (5 ng/ml) at different time points (0, 2, 8, 24, 48 h). Tubulin was used as the loading control. (D,F) Quantification of TCF7L2 protein expression. \*\*\*\**P*<0.0001; \*\*\**P*<0.001; \**P*<0.05; n.s., not significant by one-way ANOVA with Dunnett’s post-test; *n*=4. (G) Representative western blot analysis showing TCF7L2 and fibronectin expression levels after treatment with 5 ng/ml TGF-β1 (24 h) in PDGFRα-EGFP^+^ FAPs. Tubulin was used as the loading control. (H) Quantification of TCF7L2 protein expression. \*\**P*<0.005 by two-tailed Student’s t-test; *n*=3. (I) *Tcf7l2* mRNA expression levels were analyzed by quantitative PCR in C3H/10T1/2 MSCs after 2, 8 and 24 h of treatment with TGF-β1 (5 ng/ml). ****P<0.0001; **P<0.005 by one-way ANOVA with Dunnett’s post-test; *n*=3. (J) Representative western blot analysis showing TCF7L2 and PDGFRα expression levels after the treatment with actinomycin D at different time points (0, 2, 3, 4, 7 h). GAPDH was used as the loading control. (K,M) Heat maps showing the expression changes of several validated TCF7L2-target genes that are repressed or increased by TGF-β in lung fibroblasts (K) and cardiac fibroblasts (M).

Not only were the TCF7L2 protein levels decreased in response to TGF-β, but the relative levels of *Tcf7l2* mRNA were also diminished 2 h, 8 h and 24 h after TGF-β1 stimulation (Fig. 3I). *Tcf7l2* gene expression was also sensitive to the inhibition of mRNA synthesis with actinomycin D treatment, which suggests active *Tcf7l2* gene transcription and TCF7L2 translation in MSCs (Fig. 3J, Fig. S3C-E). The expression of PDGFRα was also very sensitive to the inhibition of mRNA synthesis with actinomycin D (Fig. 3J, Fig. S3C). Next, we evaluated whether TGF-β2 and TGF-β3 cytokines also impair TCF7L2 expression in MSCs. Similar to TGF-β1, both TGF-β2 and TGF-β3 strongly reduced TCF7L2 protein levels in MSCs (Fig. S3F,G). Thus, the three TGF-β ligands decrease the expression of TCF7L2 in PDGFRα^+^ cells. Next, to investigate whether the function of the Wnt-effector TCF7L2 TF was also altered by TGF-β, we determined, by quantitative PCR, the gene expression levels of several validated TCF7L2 target genes in response to TGF-β. We found that the expression of *Sox9, Axin2*, and *Tcf7l1* was repressed, while *Nfatc1, Lef1*, and *Tcf7* expression was increased in response to TGF-β1 (Fig. S3H). We did not find changes in *β-Catenin* or *Ccnd1* mRNA levels (Fig. S3H). Furthermore, RNA-seq analyses of TGF-β-treated idiopathic pulmonary fibrosis (IPF) lung (Scott et al., 2019) and cardiac fibroblasts (Schafer et al., 2017) show that TGF-β alters the expression of several validated target genes of the Wnt/β-Catenin/TCF7L2 pathway. *Dkk1, Dkk3, Lrp5, Lef1, Tcf7l1, Tgfbr2, Ctbp2, Sox9, Cxcl12, Nfatc4* expression is reduced, whereas *Fn1, Col1a1, Ctgf, Tgfb1, Foxo1, Wnt5a, Wnt5b, Wnt11, Wnt9a, Myc*, and *Oat* TCF7L2-target genes are increased after TGF-β treatment (Fig. 3K,M). Altogether, these data suggest that TGF-β signaling reduces the expression of TCF7L2, and therefore, alters its TF function in PDGFRα^+^ cells, such as FAPs, MSCs, and fibroblasts. Since TGF-β-induced myofibroblast differentiation of MSCs correlates with reduced TCF7L2 expression and impaired function, we investigated the impact that the *in vitro* differentiation towards other lineages, such as adipocytes and osteocytes, might have on *Tcf7l2* expression. We used the expression of *Adipoq* and *Runx2* as markers of effective adipogenic and osteogenic commitment in MEFs (Fig. S4). Adipogenic differentiation did not alter *Tcf7l2* mRNA expression (Chen et al., 2018), while increased expression of *Tcf7l2* was found after osteogenic differentiation (Fig. S4A). Interestingly, expression of *Tcf7l1, Lef1* and *Tcf7* also varied in response to adipogenic or osteogenic differentiation (Fig. S4A). These data suggest that different molecular mechanisms are involved in the regulation of *Tcf7l2* gene expression during differentiation to diverse cell lineages.

### Extracellular TGF-β reduces the expression of TCF7L2 in a TGF-β receptor type-I-dependent manner

Having detected a drop in the total levels of TCF7L2 TF, we next determined the extent of decrease of TCF7L2 protein levels in the nuclei of TGF-β-treated cells. A subcellular fractionation method (see Materials and Methods section for details) allowed us to determine that TCF7L2 TF was relatively abundant in the nucleus, although TCF7L2 protein was also detected in cytoplasm (Fig. S4B,C). TGF-β stimulation downregulated the expression of TCF7L2 in the nuclear and cytoplasmic fractions (Fig. 4A, Fig. S4B,C). Next, we studied further the association between TGF-β-mediated fibroblast activation and reduced TCF7L2 levels. Using confocal microscopy, we confirmed that TCF7L2 nuclear expression was decreased in response to TGF-β and that TCF7L2 repression correlated with TGF-β-induced myofibroblast phenotype in FAPs and NIH-3T3 fibroblasts (Fig. 4B,C, Fig. S4D). In addition, TGF-β signaling did not alter the subcellular distribution of TCF7L2 protein (Fig. 4D,E). Overall, these data indicate that TGF-β, in addition to inducing the differentiation of MSCs and fibroblasts into myofibroblasts, also represses the expression of the Wnt-effector TCF7L2. Then, to investigate the role of TGF-β receptors in the regulation of TCF7L2 by TGF-β, we used the TGFBR1 specific small-molecule inhibitor SB525334 (Callahan et al., 2002). TGF-βs ligand binding to the type II receptor TGFBR2 leads to recruitment and phosphorylation of the type I receptor ALK-5 (TGFBR1). Treatment with the TGFBR1 inhibitor completely abolished the downregulation of TCF7L2 expression by TGF-β1 in wild-type PDGFRα^+^ FAPs and C3H/10T1/2 MSCs, without affecting the total levels of the TCF7L2-binding partner β-catenin (Fig. 4F,G). These results suggest that TGF-β-mediated TCF7L2 downregulation requires the activation of TGF-β receptors signaling cascade.

**Fig. 4.**
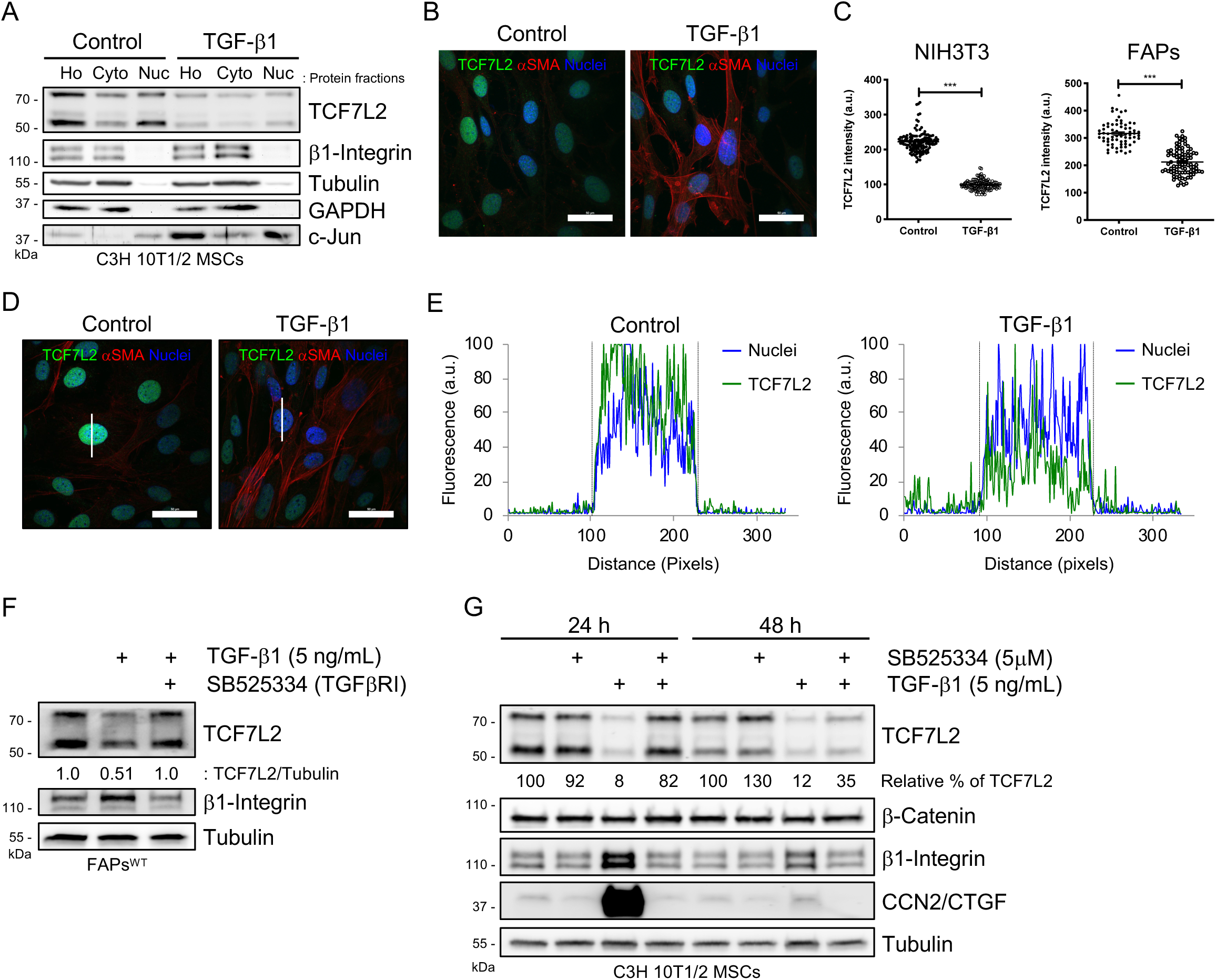
Extracellular TGF-β reduces the expression of TCF7L2 TF through TGFβR1 activation. (A) Representative western blot analysis showing TCF7L2, β1-Integrin, tubulin, GAPDH, and c-Jun expression levels in control and TGF-β1-treated C3H/10T1/2 cells. Ho: Whole cell lysate; Cyto: Cytoplasmic lysate; Nuc: Nuclei lysate. (B) Z-stack confocal images showing localization of TCF7L2 (*green*) and αSMA (*red*) in control and TGF-β1-treated (36 h) C3H/10T1/2 MSCs. Scale bars: 50 μm. (C) Quantification of the TCF7L2 fluorescence intensity in NIH-3T3 and EGFP^+^ FAPs. (a.u.: arbitrary units). Each dot represents a single cell quantified were a ROI area was previously defined. \*\*\**P*<0.001; by two-tailed Student’s t-test. *n*=3. (D) Z-stack confocal images showing localization of TCF7L2 (*green*) and αSMA (*red*) in control and TGF-β1-treated (24 h) C3H/10T1/2 MSCs. Scale bars: 50 μm. (E) Label-distribution graph showing the fluorescence intensity of TCF7L2 and Hoechst along the cell axis as shown in (D) with the *white* line. Distance is shown in pixels. Dotted lines show the nucleus-cytoplasm boundary. (a.u.: arbitrary units). (F) Representative western blot analysis showing TCF7L2 and β1-Integrin expression levels in wild-type PDGFRα^+^ FAPs co-treated for 24 h with TGFBR1 inhibitor SB525334 (5 μM) and TGF-β1 (5 ng/ml). Tubulin was used as the loading control. (G) Representative western blot analysis showing TCF7L2, β-Catenin, β1-Integrin, and CCN2/CTGF expression levels in C3H/10T1/2 MSCs co-treated for 24 h or 48 h with TGFBR1/ALK5 inhibitor SB525334 and TGF-β1. Tubulin was used as the loading control.

### TGF-β requires HDACs activity to repress *Tcf7l2* expression

The TGF-β family acts via Smad and non-Smad signaling pathways to regulate several cellular responses (Derynck and Budi, 2019; Zhang, 2017). To investigate the role of Smad and non-Smad cascades in the regulation of TCF7L2 by TGF-β, we used specific small-molecule inhibitors of Smad3, p38 MAPK, JNK, and ERK1/2 (Fig. 5A). None of these inhibitors were able to abolish the effect of TGF-β1 on the expression of TCF7L2 (Fig. 5A,B). Although we recently found that p38 MAPK participates in TGF-β-mediated downregulation of PDGFRα expression (Contreras et al., 2019c), we did not detect any significant effect of co-treatment with the p38 MAPK inhibitor SB203580 on TGF-β-induced TCF7L2 downregulation (Fig. 5A,B, Fig. S5A). Interestingly, we found that co-treatment with SIS3, a Smad3 inhibitor, augmented the inhibitory effect of TGF-β on the expression of TCF7L2 (Fig. 5A,B, Fig. S5B). Overall, these results suggest that another, not yet identified, Smad-independent pathway may participate in TGF-β-mediated downregulation of TCF7L2 expression or that rather than Smad3 activation, Smad2/Smad4 cofactors could be participating in TGF-β-mediated TCF7L2 downregulation.

**Fig. 5.**
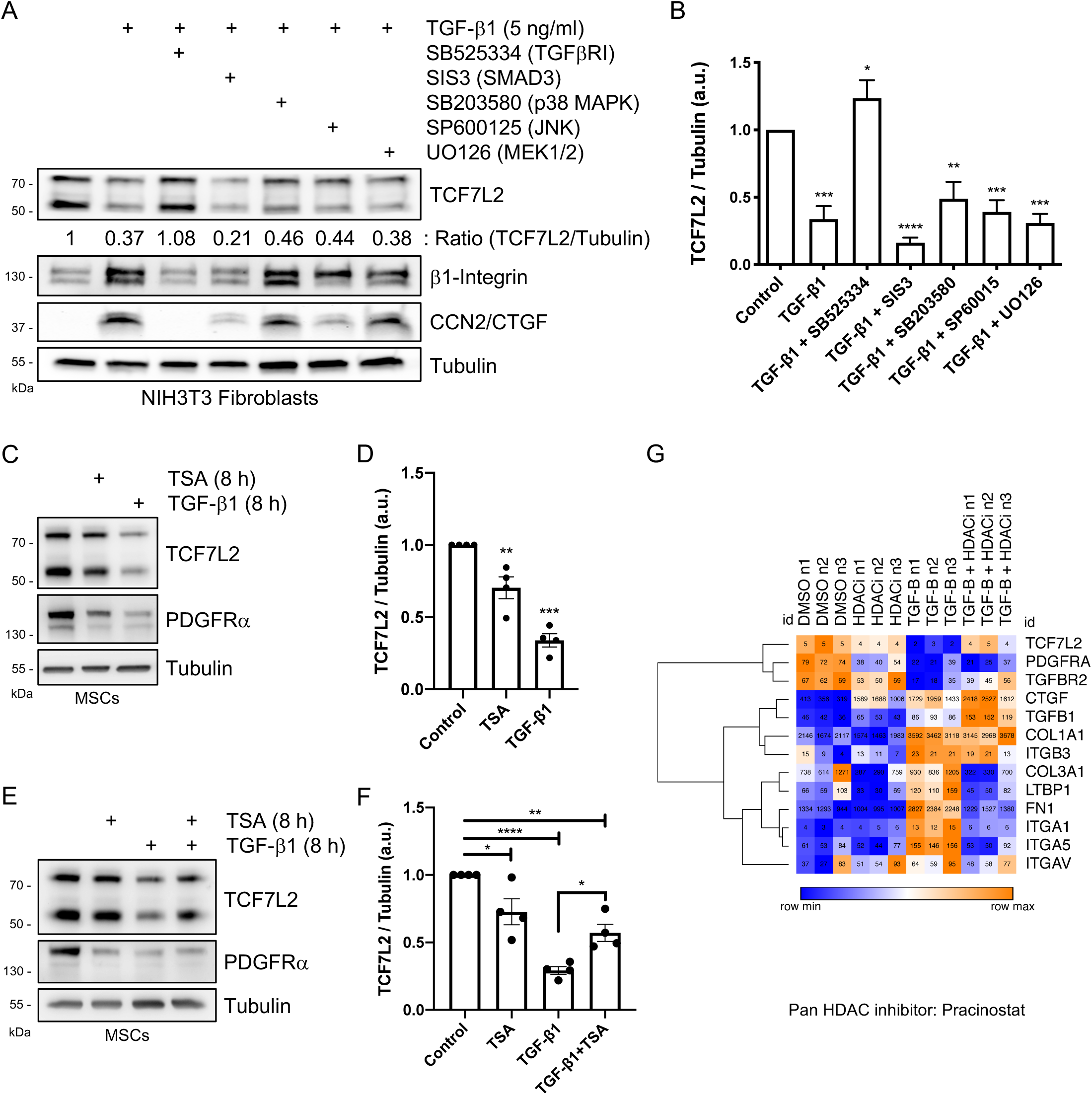
Histone deacetylases participate in TGF-β-mediated repression of TCF7L2. (A) Representative western blot analysis showing TCF7L2, β1-Integrin, and CCN2/CTGF protein levels after TGF-β1 and SB525334, SIS3, SB203580, SP600125, and UO126 pharmacological co-treatments. Tubulin was used as the loading control. (B) Quantification of TCF7L2 protein expression. \*\*\*\**P*<0.0001; \*\*\**P*<0.001; \*\**P*<0.005; \**P*<0.05; by one-way ANOVA with Dunnett’s post-test; *n*=5. (C) Representative western blot analysis showing TCF7L2 and PDGFRα protein levels after TGF-β1 (5 ng/ml) and Trichostatin A (TSA) (10 μM) treatments (8 h). Tubulin was used as the loading control. (D) Quantification of TCF7L2 protein expression. \*\*\**P*<0.001; \*\**P*<0.005 by one-way ANOVA with Dunnett’s post-test; *n*=3. (E) Representative western blot analysis showing TCF7L2 and PDGFRα protein levels after TGF-β1 and Trichostatin A (TSA) co-treatments (8 h). Tubulin was used as the loading control. (F) Quantification of TCF7L2 protein expression. \*\*\*\**P*<0.0001; \*\**P*<0.005; \**P*<0.05 by one-way ANOVA with Dunnett’s post-test; *n*=3. (G) Heat map showing *Tcf7l2* expression (RPKMs) significantly repressed by TGF-β and reversed by the pan HDAC inhibitor pracinostat. Known ECM pro-fibrotic mediators (e.g. *Collagens, Ctgf/ccn2, Fibronectin*, and *Integrins*) that are significantly increased by TGFβ and reversed by pracinostat are also shown. Each row is normalized to itself. Each column, per treatment condition, represents an individual IPF lung fibroblast donor (*n*=3).

It has been suggested that TCF7L2 might be a non-histone target of histone deacetylases (HDACs). This was based on the observation that treatment with trichostatin A (TSA), a commonly used histone deacetylase inhibitor (HDI), reduced the protein expression of TCF7L2 by 50% in the human colon cancer cell type HCT116 (Götze et al., 2014). Since HDACs regulate transcription of many genes (Bolden et al., 2006; Greet et al., 2015; Seto and Yoshida, 2014), we first evaluated whether the HDI TSA affects TCF7L2 protein levels in MSCs. Indeed, TSA treatment reduced TCF7L2 levels, although to a lesser extent than TGF-β (Fig. 5C,D). Based on recent findings that demonstrated that TGFβ-mediated fibroblast activation requires HDAC-mediated transcriptional repression (Jones et al., 2019), we examined whether inhibiting HDACs with TSA may modify TGF-β-mediated reduction of TCF7L2 expression. Therefore, the cells were treated with TGF-β1 and/or TSA for 8 h and then TCF7L2 protein levels were evaluated by western blot. Interestingly, whereas TGF-β1 decreased TCF7L2 expression, TSA partially blocked TGF-β-mediated repression of TCF7L2 expression (Fig. 5E,F). RNA sequencing analysis from recent work on IPF lung fibroblasts helped us to corroborate our results (Fig. 5G) (Jones et al., 2019). The pan HDAC inhibitor, pracinostat, attenuated TGF-β-mediated repression of *Tcf7l2* gene expression and induced extracellular matrix gene signature (Fig. 5G). Furthermore, TGF-β also impairs TCF7L2-mediated target gene expression and Wnt signaling via HDACs (Fig. S6). Taken together, these data suggest that HDACs participate in the regulation of TGF-β-mediated *Tcf7l2* repression.

### TGF-β reduces TCF7L2 protein levels by stimulating its degradation via the ubiquitin-proteasome system

To gain knowledge of the mechanisms involved in TGF-β-induced TCF7L2 repression, we performed a time-course analysis with the protein synthesis inhibitor cycloheximide. Thus, we determined that the half-life of TCF7L2 TF is quite short, being approximately 3.5 h in both proliferating C3H/10T1/2 MSCs and PDGFRα^+^ FAPs (T_1/2_=3.5 h) (Fig. 6A-C). Then, to study the molecular mechanism governing TGF-β-mediated TCF7L2 downregulation, we evaluated the human TCF7L2 interactome using BioGRID datasets (Stark et al., 2006). Remarkably, several protein-quality-control-related proteins are interacting partners of TCF7L2, among them: RNF4, RNF43, RNF138, NLK, UHRF2, UBE2I, UBE2L6, USP4, UBR5, and XIAP (Fig. 6D). Since these TCF7L2 interacting proteins belong to or are related to the ubiquitin-proteasome system (UPS), we performed *in silico* analyses and identified several top-ranked potential ubiquitination sites along the TCF7L2 protein sequence (Fig. 6E). Next, we used MG132, a potent proteasome inhibitor that reduces the degradation of ubiquitin-conjugated proteins (Nalepa et al., 2006), to evaluate the participation of the UPS in the downregulation of TCFL72 induced by TGF-β. Mechanistically, MG132 completely blocked TGF-β-mediated downregulation of TCF7L2 protein after 9 h of co-treatment in MSCs and FAPs (Fig. 6F,G, Fig. S7A,B). MG132 also increased TCF7L2 basal protein levels when administered alone, which suggests that an intrinsic ubiquitin-mediated protein degradation mechanism controls TCF7L2 steady-state levels or proteostasis (Fig. 6F,G, Fig. S7). Remarkably, a 12% SDS-PAGE gel allowed us to identify an unknown higher molecular weight form (∼65-70 kDa) of TCF7L2 not present at steady-state, but sensitive to proteasome-mediated proteolysis inhibition by MG132 (Fig. S7B). Finally, because our BioGRID analysis suggests the interaction of TCF7L2 with deubiquitinating enzymes (e.g. USP4), we evaluated the amount of TCF7L2 protein in C3H/10T1/2 MSCs treated with the ubiquitin-specific protease 7 (USP7) inhibitor HBX 41108 (HBX) in proliferating conditions (Colland et al., 2009; de la Vega, et al., 2020; Yuan et al., 2018). Treatment with only HBX diminished TCF7L2 protein levels in a time-dependent manner (Fig. 6H). We also found that USP7-mediated TCF7L2 degradation was sensitive to the inhibition of protein synthesis (Fig. 6H). Taken together, these results suggest the participation of the ubiquitin-proteasome system and ubiquitin-specific proteases in the regulation of TCF7L2 steady-state proteostasis and TGF-β-mediated repression of TCF7L2 protein expression.

**Fig. 6.**
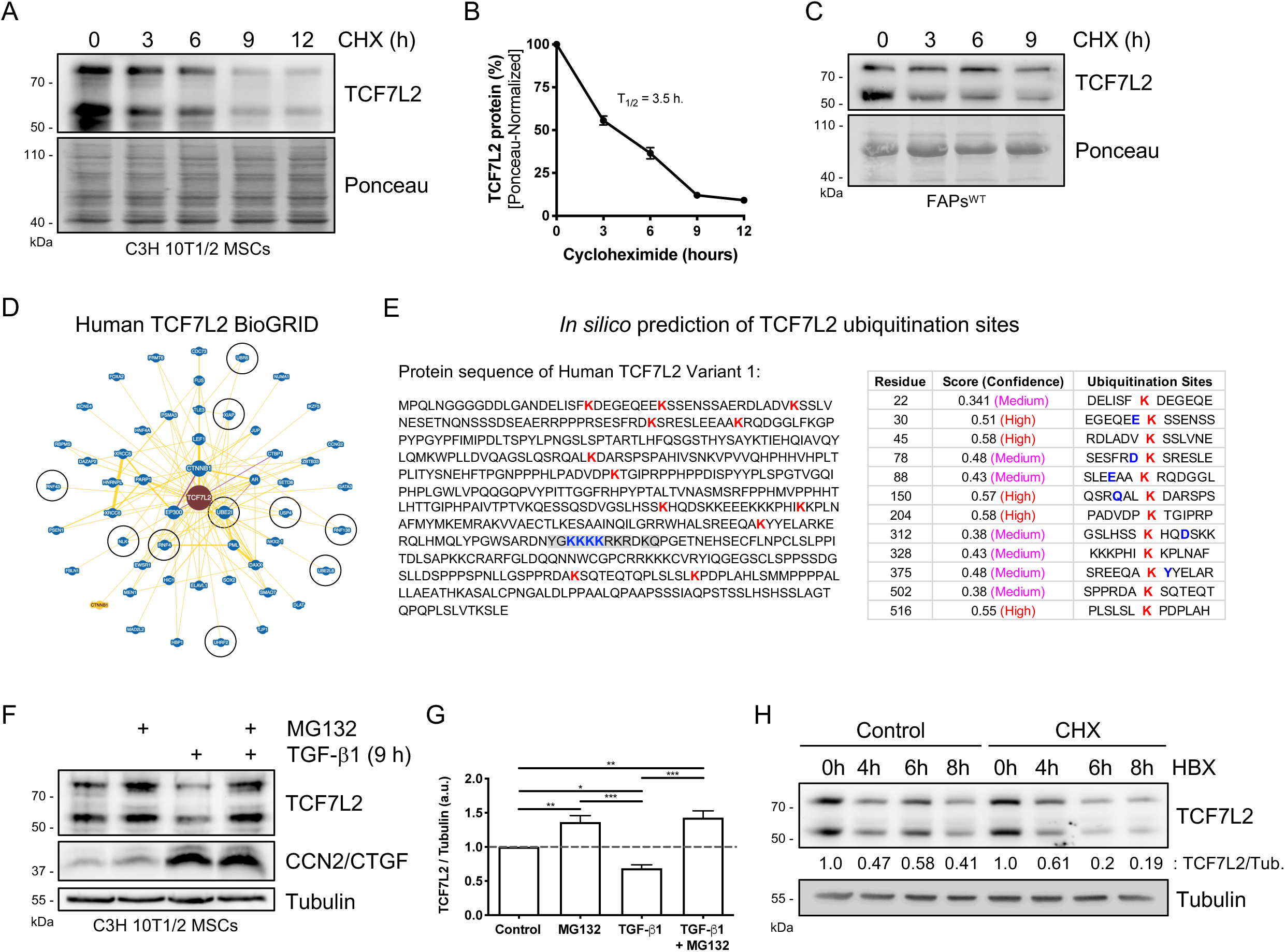
TGF-β impairs TCF7L2 protein stability via the ubiquitin-proteasome system. (A) Representative western blot analysis showing TCF7L2 protein levels after treatment with cycloheximide (CHX) (30 μg/ml) at different time points (0, 3, 6, 9, 12 h) in C3H/10T1/2 cells. Ponceau was used as the loading control. (B) Quantification of three independent experiments showing TCF7L2 protein levels (as a percentage [%]) after CHX treatment. (C) Representative western blot analysis showing TCF7L2 protein levels after treatment with CHX at different time points (0, 3, 6, 9 h) in wild-type muscle PDGFRα^+^ FAPs. (D) (A) BioGRID interactome analysis of the human TCF7L2 protein. Black circles mark the protein-protein interaction between TCF7L2 and RNF4, RNF43, RNF138, NLK, UHRF2, UBE2I, UBE2L6, USP4, UBR5, and XIAP. (E) *In silico* prediction of TCF7L2 ubiquitination sites, showing the aminoacidic sequence of human TCF7L2 protein variant 1. Potential TCF7L2-ubiquitinated lysine residues were ranked and are shown in *red*. http://www.ubpred.org; http://bdmpub.biocuckoo.org/ (F) Representative western blot analysis showing TCF7L2 and CCN2/CTGF protein levels after TGF-β1 (1 ng/ml) and MG132 (15 μM) treatments (9 h). Tubulin was used as the loading control. (G) Quantification of TCF7L2 protein levels. \*\*\**P*<0.001; \*\**P*<0.005; \**P*<0.05 by one-way ANOVA with Dunnett’s post-test; *n*=6. (H) Representative western blot analysis from three independent experiments that evaluate total levels of TCF7L2 following USP7 small-molecule inhibitor HBX 41108 (10 μM) and CHX treatments at different time points (0, 4, 6, 8 h) in C3H/10T1/2 MSCs. Tubulin was used as the loading control.

### TGF-β does not affect the expression of TCF7L2 in C2C12 myoblasts

Myoblasts are known to respond to TGF-β signaling (Droguett et al., 2006; Massagué et al., 1986; Riquelme et al., 2001; Riquelme-Guzmán et al., 2018; Schabort et al., 2009). Therefore, we investigated whether TCF7L2 downregulation by TGF-β signaling was stromal cell-type-specific or whether it also occurred in myoblasts. C2C12 myoblasts express very low levels of the TCF7L2 transcription factor and β-catenin compared to C3H/10T1/2 MSCs (Fig. 7A,B, Fig. S8A,B). Similar to what we found in stromal cells, the TCF7L2 protein was expressed predominantly in myoblasts’ nuclei (Fig. S8A). However, TGF-β1 stimulation did not alter the expression of TCF7L2, both at the protein and mRNA levels, in C2C12 myoblasts at the different concentrations used for 24 or 48 h (Fig. 7C-E, Fig. S8C,D). However, myoblasts did respond to TGF-β by increasing the expression of the two ECM components fibronectin and the matricellular protein CCN2/CTGF (Fig. 7C,E, Fig. S8C), as previously reported (Riquelme-Guzmán et al., 2018). Finally, we did not find changes in TCF7L2 subcellular distribution in myoblasts after TGF-β treatment (Fig. 7F,G, Fig. S9E). Thus, although TGF-β induces myoblast activation it does not change the expression of the TCF7L2 TF, which is expressed a relatively low levels in these cells compared to PDGFRα^+^ fibroblasts.

**Fig. 7.**
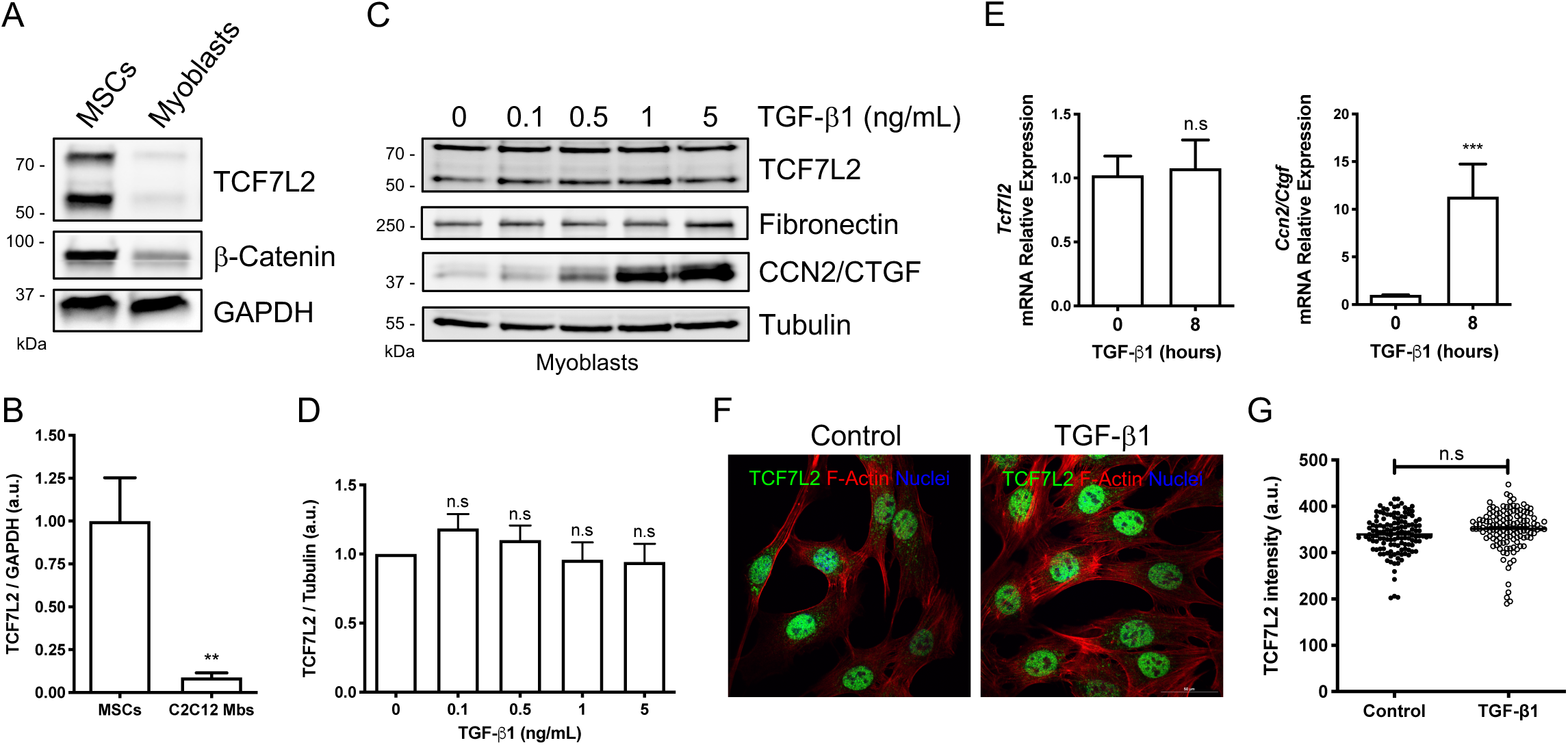
The expression of TCF7L2 remains unchanged in response to TGF-β in myoblasts. (A) Representative western blot analysis from three independent experiments, evaluating TCF7L2 and β-Catenin protein levels in proliferating C3H/10T1/2 MSCs and C2C12 myoblasts. GAPDH was used as the loading control. (B) Quantification of TCF7L2 protein levels. \*\**P*<0.005 by two-tailed Student’s t-test. *n*=3. (C) Representative western blot analysis showing TCF7L2, fibronectin, and CCN2/CTGF protein levels after treatment with different concentrations of TGF-β1 for 24 h. (D) Quantification of TCF7L2 protein levels. n.s, not significant by two-tailed Student’s t-test. *n*=3. (E) *Tcf7l2* and *CCN2/CTGF* mRNA expression levels were analyzed by quantitative PCR in C2C12 myoblasts after 8 h of treatment with TGF-β1 (5 ng/ml). \*\*\**P*<0.001; n.s, not significant by two-tailed Student’s t-test. *n*=3. (F) Representative Z-stack confocal images showing localization of TCF7L2 (*green*) and F-actin (*red*) in control and TGF-β1-treated (24 h) C2C12 myoblasts. Scale bar: 50 μm. (G) Quantification of the TCF7L2 fluorescence intensity in C2C12 myoblasts. Each dot represents a single cell quantified were a ROI area was previously defined. n.s; not significant by two-tailed Student’s t-test. *n*=3. (a.u.: arbitrary units).

## DISCUSSION

Here, we first reported differential gene expression of the four Wnt Tcf/Lef TFs in PDGFRα^+^ fibroblasts from skeletal muscle and cardiac tissue, MSCs cell lines, and MEFs. Second, we established that TCF7L2 is a novel TGF-β target gene. Therefore, not only the gene and protein expression of TCF7L2 are strongly reduced by TGF-β signaling but also the expression of several TCF7L2-target genes is altered upon TGF-β stimulation (Fig. 8). Third, we showed via TCF7L2 BioGRID-based interaction network, *in silico* prediction of TC7L2 ubiquitination residues, and proteasome inhibition, that TGF-β regulates TCF7L2 protein stability via the UPS (Fig. 8). We also described that two pan HDACs inhibitors, TSA and pracinostat, counteracted TGF-β-mediated repression of TCF7L2 (Fig. 8). Finally, we observed that TGF-β-driven TCF7L2 downregulation is fibroblast-specific, as this effect did not occur in C2C12 myoblasts.

**Fig. 8.**
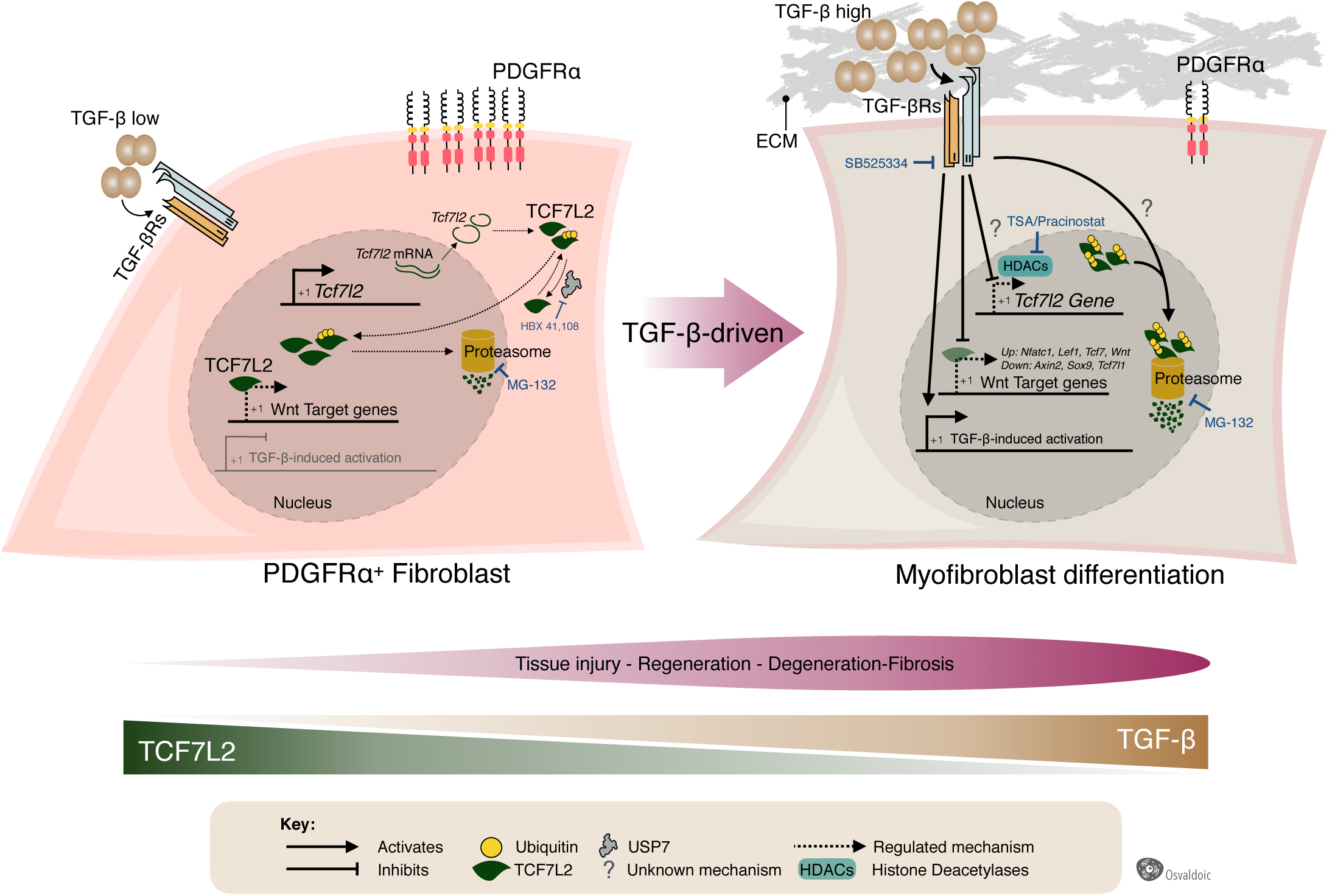
Model of TGF-β-driven fibroblast-myofibroblast differentiation and the Wnt-TCF7L2 regulatory network. MSCs and fibroblasts express high levels of TCF7L2 in the resting-quiescent state, but they lose this TF post-activation with TGF-β. *Tcf7l2* gene expression is active in unstimulated cells, and therefore, it continuously produces the TCF7L2 protein. TCF7L2 localizes mainly in the cell nucleus where it recognizes its target genes to activate or repress their expression, depending on the cell context and its transcriptional partners. The ubiquitin-proteasome system constantly regulates the proteostasis of TCF7L2. Following muscle damage, TGF-β ligands released from macrophages (paracrine) and FAPs (autocrine) bind to TGF-βRs. This event activates TGFBRs-dependent signalling cascades that downregulate TCF7L2 expression and impair TCF7L2-dependent gene expression and Wnt/β-catenin gene signature. Mechanistically, inhibition of the 26S proteasome activity with MG132 blocks TGF-β-mediated TCF7L2 protein degradation. Also, HDACs pan inhibitors (TSA and Pracinostat) attenuate TGF-β-mediated repression of TCF7L2 expression and ECM gene activation.

Tcf/Lef can act as activators or repressors of gene transcription, like other high-mobility group box-containing proteins, mostly depending on their binding cofactors, activators or repressor and in a tissue- and cell-specific fashion (Frietze et al., 2012; Jin, 2016; Lien and Fuchs, 2014; Tang et al., 2008; Weise et al., 2010). TCF7L2, emerges as an interesting TF to study because TCF7L2^+^ MSCs are increased in fibrotic muscles from the *mdx* mice and DMD (Contreras et al., 2016; Pessina et al., 2015), and also in damaged and atrophied muscles of the symptomatic amyotrophic lateral sclerosis (ALS) transgenic mice hSOD^G93A^ (Gonzalez et al., 2017). TCF7L2^+^ cells peak at day 5 following acute injury and then return to their basal levels once the damage is resolved (Murphy et al., 2011). In addition, TCF7L2 is a known transcriptional regulator in several cancers (Clevers, 2006; Cosin-Roger, 2019; Jin, 2016; Ravindranath et al., 2008; Tang et al., 2008; van de Wetering et al., 2002), and is the gene more closely associated with risk of type 2 diabetes to date (Grant et al., 2006; Jin, 2016). Nevertheless, most of the knowledge concerning canonical Wnt signaling comes from β-Catenin translocation to the nucleus or dysregulated activity of upstream β-Catenin-stability regulators like GSK-3β or CSNK1, but the precise contribution of Tcf/Lef members to fibrogenesis has not been addressed yet. Even though TCF7L2 has been described as a marker for muscle CT fibroblasts (Contreras et al., 2016; Kardon et al., 2003; Mathew et al., 2011; Merrell et al., 2015), increasing evidence suggests that these cells share many properties with muscle PDGFRα^+^ FAPs (Contreras et al., 2019b; Malecova et al., 2018; Wosczyna and Rando, 2018).

A recent report demonstrated that TCF7L2 protein but not mRNA levels are increased during adipocyte differentiation, thereby TCF7L2 plays a regulatory role in fibroblast-adipocyte fate decisions *in vitro* and *in vivo* (Chen et al., 2018). In agreement with Chen and Cols. (2018) we did not find *Tcf7l2* gene expression changes after adipogenic differentiation. However, the precise role of TCF7L2 in regulating the fate and lineage restriction of fibroblast still needs to be addressed. It is important to mention that our work may have some experimental limitations. For example, we have not explored the role of different Tcf/Lef TFs on the fate of MSCs yet. Here, we also reported that *Tcf7l1* (*Tcf3*) is a Tcf/Lef gene with high expression in PDGFRα^+^ cells. The regulation of the expression of *Tcf7l1* seems to resemble that of *Tcf7l2* but its role in fibroblast biology still needs to be further addressed. Interestingly, *Tcf7* and *Lef1* were expressed at very low levels under resting conditions, but they were strongly upregulated by tissue damage and TGF-β. This might be explained as a compensatory regulation of them or as a TCF7L2-mediated response. Tcf7 and Lef1 are target genes of TCF7L2 (Frietze et al., 2012; Lien et al., 2014), which may explain why these two members are upregulated when TCF7L2 is downregulated by TGF-β. However, future studies need to address the questions raised above.

The UPS has emerged as an essential component in cell and molecular biology via regulation of cellular proteostasis in homeostasis and disease (Nalepa et al., 2006; Yuan et al., 2018). Here, we demonstrated the participation of both the UPS and the deubiquitinase enzyme USP7 in the regulation of TCF7L2 steady-state levels and stability in response to TGF-β. Although we did not address the ubiquitination status of TCF7L2, to our knowledge this is the first study suggesting potential cross-modulation between TGF-β and the Wnt pathway through deubiquitinating enzymes and the UPS. Post-translational modifications of TCF7L2 occur at the phosphorylation (Ishitani et al., 2003), acetylation (Elfert et al., 2013), and SUMOylation level (Yamamoto et al., 2003). Although recent reports demonstrated the ubiquitination of some Tcf/Lef members (Han et al., 2017; Song et al., 2018, Yamada et al., 2006), the extent and importance of ubiquitin-mediated regulatory mechanisms for these TF activity are still elusive. TCF7L1 half-life was reported to be longer than 12 h, where the DNA-binding of TCF7L1 regulates its protein stability, and therefore, Wnt signaling (Shy et al., 2013). Since TGF-β signaling is a known chromatin-modifying factor (Massagué, 2012), an intriguing possibility is that TCF7L2 DNA-binding regulates its stability and that TGF-β-induced chromatin remodeling might negatively affect TCF7L2 stability. However, the complexity of different molecular events maintaining TCF7L2 protein or mRNA levels in PDGFRα^+^ cells remains unclear. A limitation of this study is that we did not investigate the role of TCF7L2 in TGF-β-mediated ECM remodeling and myofibroblast differentiation. As TCF7L2 TF is known for regulating the expression of thousands of genes in a cell-type-specific manner (Frietze et al., 2012; Schuijers et al., 2014; Tang et al., 2008; Weise et al., 2010), we propose that TGF-β-induced TCF7L2 TF downregulation should have a massive and profound impact on fibroblast’s transcriptome.

Recently, histone deacetylase inhibitors (HDACi) have emerged as potential compounds for use in pre-clinical and clinical studies to improve tissue regeneration and repair in DMD (Bettica et al., 2016; Minetti et al., 2006). Central to this idea is that PDGFRα^+^ cells are specifically targeted by HDACi in muscular dystrophies (Mozzetta et al., 2013; Saccone et al., 2014). HDACi reduce DMD progression by increasing muscle regeneration while inhibiting fibro-fatty differentiation of PDGFRα^+^ cells. Mechanistically, HDACi favor intercellular communication between PDGFRα^+^ FAPs and myogenic progenitors during regeneration and repair (Bettica et al., 2016; Mozzetta et al., 2013; Saccone et al., 2014). Here, we observed that two well-characterized and known pan-HDACi reduce TGF-β-mediated ECM gene signature and also block TGF-β-induced downregulation of *Tcf7l2* expression. This corresponds with our results showing that HDACi modulate TGF-β-mediated Wnt-TCF7L2 downstream genes. Further studies should unravel the mechanism by which HDACi regulate Wnt-TCF7L2-dependent gene network.

Canonical Wnt signaling activation through Wnt-1 upregulates collagen ECM deposition and promotes fibroblasts differentiation into myofibroblasts (Akhmetshina et al., 2012). Similarly, TGF-β activates the canonical Wnt cascade by inducing β-Catenin nuclear accumulation and increasing Tcf/Lef-responsive elements activity in reporter assays. Therefore, TGF-β mediates the reduction of the expression of the Wnt inhibitor DKK-1 (Akhmetshina et al., 2012). This seems to be in agreement to our results showing that TGF-β reduces the expression of several TCF7L2 target genes such as *Dkk1, Dkk3, Lrp5, Lef1, Tcf7l1, Tgfbr2, Ctbp2, Sox9, Cxcl12*, and *Nfatc4* in IPF lung and heart fibroblasts. Furthermore, pharmacological inhibition of Wnt/β-catenin signaling by the small molecule ICG-001, reduced proliferation and TGF-β1-induced myofibroblast differentiation of lung resident MSCs *in vitro* and triggered organ protection via attenuation of bleomycin-induced lung fibrosis *in vivo* (Cao et al., 2018). These studies and others suggest that the canonical Wnt cascade is required for TGF-β-mediated effects, and vice versa (Cosin-Roger et al., 2019; Działo et al., 2018; Girardi and Le Grand, 2018; Piersma et al., 2015). Here, we hypothesize that extracellular TGF-β release after injury primes PDGFRα^+^ progenitor cells into differentiating TCF7L2 non-expressing myofibroblasts, which are then refractory to the self-renewal Wnt signals. Central to this idea is the significance of restraining an exacerbated stromal response to control scar formation. Thus, Wnt/TCF7L2 and TGF-β signaling could interact in a cell-specific and complex arrangement during regeneration and repair.

Emerging studies suggest a functional crosstalk between Wnt cascade and TGF-β signaling in modulating MSCs activation and fate (Burgy and Königshoff, 2018; Cosin-Roger et al., 2019; Vallée et al., 2017; Hamburg-Shields, et al., 2015). For example, the canonical Wnt3a ligand upregulates TGF-β signaling via Smad2 in a β-catenin-dependent mechanism, and therefore, promotes the differentiation of fibroblasts into myofibroblasts (Carthy et al., 2011). Moreover, Wnt/β-catenin/TCF7L2 pathway mediates the transcriptional regulation of the TGF-β-target gene, TMEPAI (Nakano et al., 2010). TGF-β promotes the secretion of Wnt proteins, via transforming growth factor beta-activated kinase 1 activation, which in turn activates the canonical Wnt cascade (Blyszczuk et al., 2016). Therefore, this TGF-β-dependent Wnt secretion induces myofibroblasts formation and myocardial fibrosis progression (Blyszczuk et al., 2016). Elevated canonical Wnt/β-catenin signaling is found in dystrophic *mdx* muscles and controls MuSCs fate via crosstalk with TGF-β2 (Biressi et al., 2014). Also, TGF-β induces the production of the canonical ligand Wnt3a, which in turn increases TGF-β secretion, establishing an amplification circuit between TGF-β and Wnt signaling pathways in cardiac fibroblasts (Seo et al., 2019). Also, it has been demonstrated that TGF-β3-induced epithelial-mesenchymal transformation proceeds via transcription complexes of Smad2/Smad4/LEF1 that directly inhibit E-cadherin gene expression (Nashaw et al., 2007). Tian et al., (2013) reported that β-catenin is a co-factor of Smad3 during TGF-β1-mediated epithelial-mesenchymal transformation. Therefore, these studies and ours suggest a novel interplay between TGF-β and the canonical Wnt pathway.

The study of the heterogeneity and plasticity of tissue-resident mesenchymal stromal populations emerges as an attractive field to understand regeneration versus degenerative fibrosis (Riquelme-Guzmán and Contreras, 2020; Mahmoudi et al., 2019; Lemos and Duffield, 2018; Lynch and Watt, 2018). Different MSC populations and their lineage may have intrinsic properties that favor either permanent scar formation or regeneration via scar regression (Driskell et al., 2013; Malecova et al., 2018; Plikus et al., 2017; Rinkevich et al., 2015; Rognoni et al., 2018; Soliman et al., 2020; Furtado et al., 2016). Thus, investigating different sub-populations of fibroblasts, with particular niches and genetic programs, is important to understand how these cells and their progeny influence wound healing, tissue repair, and regeneration. Dynamic downregulation of fibroblast markers following damage has also been identified in resident PDGFRα^+^ cardiac fibroblasts (Asli et al., 2018 preprint; Farbehi et al., 2019; Kanisicak et al., 2016; Tallquist and Molkentin, 2017; Soliman et al., 2020) and skeletal muscle (Contreras et al., 2019c; Malecova et al., 2018). We have recently reported that the *in vivo* and *in vitro* expression of PDGFRα is strongly downregulated by damage-associated TGF-β signaling in skeletal muscle and heart PDGFRα^+^ fibroblasts (Contreras et al., 2019c). Conversely, TGF-β mediates the differentiation of PDGFRα^+^ cells into myofibroblasts at the expenses of their adipogenic differentiation (Contreras et al., 2019c; Uezumi et al., 2014a; Uezumi et al., 2011). In summary, the work of others and our results suggest that stromal stem cell and/or progenitor markers are often downregulated by TGF-β, probably after a complex array of niche signals during regeneration or disease, as cells change phenotypically towards an activated cell state. These mesenchymal progenitor and/or fibroblast downregulated markers include Sca-1, PDGFRα, Tcf21, and Hic1 (Asli et al., 2018 preprint; Contreras, 2019d; Contreras et al., 2019c; Fu et al., 2018; Kanisicak et al., 2016; Scott et al., 2019; Soliman et al., 2020). With our work, we added TCF7L2 to the growing list.

Our understanding of the cellular and molecular determinants of fibrosis has advanced immensely (Ceco and McNally, 2013; Kim et al., 2018; Piersma et al., 2015; Smith and Barton, 2018). Nevertheless, the lack of successful antifibrotic therapy to date prompts us to continue the search for new potential candidates to fight against non-malignant proliferative disorders. Compelling evidence from this work raises the idea that TCF7L2 should be explored as a therapeutic target in fibrotic diseases, as the Wnt β/Catenin pathway emerges as a novel and attractive signaling cascade to target for improving tissue function in myopathies, fibrosis-related disorders, and aging.

## MATERIALS AND METHODS

### Mice and study approval

Housing, husbandry and experimental protocols were conducted in strict accordance and with the formal approval of the Animal Ethics Committee of the Pontificia Universidad Católica de Chile (Doctoral ID protocol: 160512005) and following institutional and national guidelines at the University of British Columbia, Canada. Mice were housed in standard cages under 12-h light-dark cycles and fed *ad libitum* with a standard chow diet. Five-month-old C57BL/10ScScJ male mice (hereafter referred to as wild type, WT; stock #000476) and dystrophic C57BL/10ScSn-Dmdmdx/J mice (stock #001801) male mice (both from Jackson Laboratories) were used in experiments for Fig. 1E, Fig. 2, and Fig. S3H. *Pdgfra*^*tm11(EGFP)Sor*^ mice (hereafter referred to as PDGFRα^H2BEGFP^ mice) were purchased from Jackson Laboratories (stock #007669 B6.129S4-Pdgfra^tm11(EGFP)Sor^/J; Hamilton et al., 2003). For FAP detection in *mdx* muscles, we crossed male C57Bl/10ScSn-*mdx* mice with hemizygous female B6.129S4-Pdgfra^tm11(EGFP)Sor^/J mice. We used the F1 male *mdx*;PDGFRα^H2BEGFP^ offspring (5- to 6-month-old), and the comparisons were performed among siblings. All surgeries were performed after the mice had been anesthetized with 2.5–3% of isoflurane gas in pure oxygen. The mice were euthanized with cervical dislocation at the ages indicated in each figure, and the tissues were immediately processed, either by direct freezing in liquid nitrogen for protein and RNA extraction or in 2-methyl butane cooled with liquid nitrogen for histological analysis as described below.

### Muscle acute injury

For acute glycerol injury, the tibialis anterior (TA) muscle from 2- to 3-monthold PDGFRα^H2BEGFP^ mice was injected with 50 μl 50% v/v glycerol. Tissue collection was performed as indicated in the corresponding figures after glycerol injections. Notexin muscle damage was induced by intramuscular injection of 0.15 μg notexin snake venom (Latoxan) into the TA muscle (Contreras et al., 2019c; Joe et al., 2010; Lemos et al., 2015). Non-injected muscles from the contralateral limb were used as controls. Muscles were isolated and collected for analysis at the indicated time points in the corresponding figures.

### Tissue preparation, flow cytometry and FACS

Tissue preparation for skeletal muscle and heart FAPs was performed mainly as described before (Contreras et al., 2019c; Soliman et al., 2020). One-step digestion of tissue for FAPs was performed mainly as described before with some modifications (Judson et al., 2017; Lemos et al., 2015). All the steps were performed on ice unless otherwise specified. Briefly, skeletal muscle from both hindlimbs (limb FAPs), and diaphragm (diaphragm FAPs) was carefully dissected, washed with 1×PBS, cut into small pieces with scissors until homogeneous. Collagenase D (Roche Biochemicals) 1.5 U/ml and Dispase II (Roche Biochemicals) 2.4 U/ml, in 2.5 mM CaCl2, was added to every two hindlimbs in a total volume of 3 ml per mouse, and the preparation was placed at 37°C for 45 min with rotation. Preparations were passed through a 70 μm, and then 40 μm cell strainer (Becton Dickenson), and washed with FACS buffer (PBS, 2% FBS, 2 mM EDTA pH 7.9). Resulting single cells were collected by centrifugation at 1000 *g* for 5–10 min. Cell preparations were incubated with primary antibodies for 20–30 min at 4°C in FACS buffer at ∼3×107 cells/ml.We used the following monoclonal primary antibodies: anti-CD31 (clones MEC13.3, Cat. no. 553372, Becton Dickenson; clone 390, Cat. no. CL8930F-3, 1:500, Cedarlane Laboratories), anti-CD45 (clone 30-F11, Cat. no. 557659, 1:400, Becton Dickenson), anti-CD45.1 (1:400; clone A20, Cat. no. 553775, 1:400, Becton Dickenson), anti-CD45.2 (clone 104, Cat. no. 11-0454-85, eBiosciences), anti-Sca-1 (1:2000–1:5000; clone D7, Cat. no. 25-5981-82, Invitrogen) and anti-α7 integrin (1:11500; Clone R2F2, Cat. no. 67-0010-05, AbLab). For all antibodies, we performed fluorescence minus one control by staining with appropriate isotype control antibodies (rat IgG2a kappa, PE-Cyanine7, clone eBR2a, Cat. no. 25-4321-82, eBioscience. 1:400; mouse anti-IgG2a k, FITC, clone G155-178, BD, Cat No: 553456; rat IgG2b kappa, APC, clone eB149/10H5, Cat. no. 17-4031-82 – all from eBioscience). To assess viability, cells were stained with propidium iodide (1 μg ml–1) and Hoechst 33342 (2.5 μg ml–1) and resuspended at ∼1×106 cells ml–1 immediately before sorting or analysis. The analysis was performed on a LSRII (Becton Dickenson) flow cytometer equipped with three lasers. Data were collected using FacsDIVA software. Cell sorting was performed on a FACS Vantage SE (Becton Dickenson), BD Influx flow cytometer (Becton Dickinson), or FACS Aria (Becton Dickenson), all equipped with three lasers, using a 100-μm nozzle at 18 psi to minimize the effects of pressure on the cells. Sorting gates were strictly defined based on isotype control (fluorescence minus one) stains. All flow cytometry data were analyzed using FlowJo 10.5.3v.

### Reagents

The TGFBR1 inhibitor SB525334 (used at 5 μM; S8822, Sigma-Aldrich), p38 MAPK SB203580 inhibitor (used at 20 μM; 5633, Cell Signaling Technology), PI3K/AKT inhibitor LY294002 (used at 10 μM; 440202, Merck-Calbiochem), the inhibitor of MEK1/2/ERK1/2 kinases UO126 (used at 10 μM; 9903, Cell Signaling Technology), Smad3 inhibitor SIS3 (used at 6 μM; 1009104-85-1, Merck-Calbiochem), the inhibitor of JNK activity SB600125 (used at 20 μM; Cell Signaling Technology), trichostatin A (TSA) (used at 10 μM; T8552, Sigma-Aldrich), and the inhibitor of USP7 activity HBX 41108 (used at 10uM; 4285; Tocris) were all diluted in DMSO. DMSO alone was used as a control. Cycloheximide (C104450, Sigma-Aldrich) was diluted in ethanol and used at 30μg/ml final concentration. All the inhibitors used were added at the same time and co-incubated with TGF-β1. Other reagents, unless otherwise is indicated, were purchased from Sigma-Aldrich.

### Cell culture and nuclei cytoplasmic fractionation

The murine mesenchymal stromal cell (MSC) cell line C3H/10T1/2, Clone 8, and the embryonic fibroblast cell line NIH-3T3 were obtained from American Type Culture Collection (ATCC) and grown at 37°C in 5% CO2 in growth medium (GM): high-glucose Dulbecco’s modified Eagle’s medium (DMEM) (Invitrogen) with 10% fetal bovine serum (FBS; Hyclone) and supplemented with antibiotics (Gutiérrez et al., 2015). The murine C2C12 myoblast cell line (American Type Culture Collection (ATCC), VA, USA) were cultured at 37 °C in 8% CO2 in growth medium (GM); DMEM high glucose (Invitrogen, CA, USA) with 10% fetal bovine serum (FBS) (Hyclone, UT, USA) and supplemented with antibiotics. Cells were treated with recombinant hTGF-β1 (#580702, Biolegend, USA), recombinant hTGF-β2 (#583301, Biolegend, USA), recombinant hTGF-β3 (#501123524, eBioscience, CA, USA) in Dulbecco’s modified Eagle’s medium (DMEM) supplemented with 2% (v/v) FBS and penicillin/streptomycin in a 5% CO2 atmosphere at concentration and time indicated in the corresponding figure legend. Adipogenic or osteogenic differentiation of MEFs was induced for 28 days with MesenCult™ Adipogenic Differentiation Kit (Mouse) (STEMCELL Technologies, Canada) and MesenCult™ Osteogenic Stimulatory Kit (Mouse), respectively. Our cell cultures were periodically tested to ensure no mycoplasma contamination using polymerase chain reaction (PCR). Cell fractionation was performed mainly according the principles of rapid, efficient and practical (REAP) method for subcellular fractionation with a few modifications (Suzuki et al., 2010). Briefly, the initially PBS-scraped and pelleted cells were resuspended, by pipetting up/down and vortexing for 5 s, using 900 μL of “cytoplasmic buffer”: 10mM Tris pH 7.5, 3mM MgCl_2_, 100mM NaCl_2_, 1mM EGTA, 0.25% (v/v) Nonidet P-40. Next, 300 μL of the initial lysate was removed and kept as “whole cell lysate” or Ho (Fig. 4A). The remaining (∼600 μL) material was centrifuged for 10 sec (two times) in 1.5 ml micro-centrifuge tubes and 300 μL of the supernatant was removed and kept as the “cytosolic fraction”. After the remaining supernatant was removed, the pellet (∼20 μL) was resuspended with 180 μL of 1 × Laemmli sample buffer and designated as “nuclear fraction”. 100 μL of 4 x Laemmli sample buffer was added to both the whole cell lysate and the cytosolic fractions. Finally, samples were sonicated and 30 μL, 30 μL and 15 μL of whole cell lysate, cytoplasmic and nuclear fractions, respectively, were loaded and electrophoresed using sodium dodecyl sulfate polyacrylamide gel electrophoresis (SDS-PAGE) (see below).

### FAP cell culture

PDGFRα^+^ FAPs were FACS-isolated from either wild-type or PDGFRα^H2BEGFP/+^ mice and grown in high-glucose DMEM (Invitrogen), supplemented with 10% FBS, 1% sodium pyruvate, and 2.5 ng/ml bFGF (Invitrogen) at a density of 15,000 cell/cm^2^ in a 48-well plate or 24-well plate. Cells were isolated from undamaged muscles. For the TGF-β1 treatment experiment, after 72 h to 96 h in culture and 70–80% confluence, FAPs were stimulated with 5 ng/ml TGF-β1 (Contreras et al., 2019c). Cells were then collected for further analyses.

### Protein extraction and western blot analysis

Protein extracts from cells were obtained using RIPA 1x lysis buffer (9806, Cell Signaling, MA, USA) plus protease/phosphatase inhibitors (#P8340/#P0044, Sigma-Aldrich, USA). Whole-muscle and heart extracts were obtained by homogenization at 4°C in RIPA 1x lysis buffer plus protease/phosphatase inhibitors. The cells were sonicated for 10 s and centrifuged at 9,000 *g*. Whole-tissue homogenates were clarified by centrifugation at 10,000 *g* for 5 min. Proteins were quantified with the Micro BCA assay kit, following the manufacturer’s instructions (Pierce, IL, USA). Protein extracts (30–60 μg) were subjected to SDS-PAGE electrophoresis in 9-10% (or 12% in Fig. S7B) polyacrylamide gels, transferred to PDVF membranes (Millipore, CA, USA), and probed with primary antibodies: goat anti-PDGFRα (1:1000; AF1062, R&D Systems), rabbit anti-TCF4/TCF7L2 (C48H11) (1:1000; 2569, Cell Signaling), rabbit anti-c-Jun (60A8) (1:1000; 9165, Cell Signaling), rabbit anti-β-Catenin (1:2000; 9562, Cell Signaling), rabbit anti-histone 3 H3 (1:1000; 9715, Cell signaling), mouse anti-alpha smooth muscle actin (αSMA) (1:2000; Cat. No A5228, Sigma-Aldrich, St. Louis, MO, USA), goat anti-CCN2/CTGF (1;500; Cat. no. sc-14939, Santa Cruz Biotechnology), rabbit anti-Integrin β1 (M-106) (1:1000; sc-8978, Santa Cruz Biotechnology), rabbit anti-fibronectin (1:2000; F3648, Sigma-Aldrich), mouse anti-Myosin Skeletal Fast (1:1000; M4276, Sigma-Aldrich), rabbit anti-myogenin (1:500; sc-576, Santa Cruz), mouse anti-GAPDH (1:5000; MAB374, Millipore), mouse anti-α-tubulin (1:5000; T5168, Sigma-Aldrich), Then, primary antibodies were detected with a secondary antibody conjugated to horseradish peroxidase: mouse anti-goat IgG (1:5000; 31400), goat anti-rabbit IgG (1:5000; 31460) and goat anti-mouse IgG (1:5000; 31430), all from Pierce. All immunoreactions were visualized by enhanced chemiluminescence Super Signal West Dura (34075, Pierce) or Super Signal West Femto (34096, Pierce) by a ChemiDoc-It HR 410 imaging system (UVP). Western blot densitometry quantification was done using Fiji software (ImageJ version 2.0.0-rc/69/1.52n). Briefly, minimum brightness thresholds were increased to remove background signal. Remaining bands were bracketed, plot profiles generated, and area under histograms auto traced. Protein levels were normalized with the levels of the loading control. Ponceau S Red Staining Solution (0.1% (w/v) Ponceau S in 5% (v/v) acetic acid) was used.

### Indirect immunofluorescence and microscopy

Cells immunofluorescence was performed as previously described (Contreras et al., 2018). For tissue section immunofluorescence, flash-frozen muscles were sectioned at 7 μm, fixed for 15 min in 4 % paraformaldehyde, and washed in phosphate-buffered saline (PBS). Cells and tissue sections were blocked for 30–60 min in 1 % bovine serum albumin (BSA) plus 1 % fish gelatin in PBS, incubated overnight at 4 °C in primary antibody: rat anti-Laminin-α2 (1:250, Cat. No L0663, Sigma-Aldrich), rabbit anti-Collagen Type I (1:250, Cat. No A34710, Abcam), rabbit anti phospho-Smad3 antibody (1:100; Cat. No 9520S, Cell Signaling), mouse anti-alpha smooth muscle actin (αSMA) (1:250, Cat. No A5228, Sigma-Aldrich, St. Louis, MO, USA). Then, washed in PBS, incubated for 1 h or at room temperature with a secondary antibody (Alexa-Fluor-568 donkey anti-rabbit IgG (H+L), Cat. no. A10042, Life Technologies; Alexa-Fluor-594 donkey anti-rabbit IgG (H+L), Cat. no. A21207, Life Technologies; Alexa-Fluor-488 goat anti-rabbit IgG (H+L), 1:500, Cat. no. A11008, Life Technologies; Alexa-Fluor-555 donkey anti-mouse IgG (H+L), Cat. no. A31570, Invitrogen; Alexa-Fluor-568 goat anti-rat IgG (H+L), Cat. no. A11077, Life Technologies; all used at 1:500) and washed in PBS. Hoechst 33342 stain (2 mg/ml) and wheat germ agglutinin (WGA) Alexa Fluor 594 conjugate (#W11262, Invitrogen, CA, USA) were added for 10 min in PBS before the slides were mounted, according to the supplier’s instructions. Slides were then washed in PBS and mounted with fluorescent mounting medium (DAKO, USA). To stain F-actin Alexa Fluor 568 Phalloidin was added to the cells according to provider’s instructions for 10 minutes (#A12380, Thermo-Fisher, MA, USA). Cells were imaged on a Nikon Eclipse C2 Si Confocal Spectral Microscope or Nikon Eclipse Ti Confocal Microscope using Nikon NIS-Elements AR software 4.00.00 (build 764) LO, 64 bit. Confocal images were acquired at the Unidad de Microscopía Avanzada (UMA), Pontificia Universidad Católica de Chile, using a Nikon Eclipse C2 Si confocal spectral microscope. Plan-Apochromat objectives were used (Nikon, VC 20× DIC N2 NA 0.75, 40× OIL DIC H NA 1.0, and, VC 60× OIL DIC NA 1.4). Cytospin confocal microscopy was performed using a Nikon eclipse Ti Confocal Microscope with a C2 laser unit. Confocal microscopy images shown in Fig. 1G, Fig. 2C and Fig. S1G were composed using maximum-intensity projection z-stack reconstructions (0.3 μm each stack) of 7-μm-thick transversal sections or cultured cells. Then, we automatically analyzed the intensity of fluorescence (amount) of TCF7L2 in TCF7L2^+^ cells using the analyze particles plugging, and manually counted the cells using the cell counter plugging from Fiji software (ImageJ version 2.0.0-rc/69/1.52n, NIH). Counts of 6–8 randomly chosen fields were averaged from four independent experiments.

### RNA isolation, reverse transcription, and quantitative real-time polymerase chain reaction (RT-qPCR)

Total RNA from cultured cells was isolated using TRIzol (Invitrogen, CA, USA) according to the manufacturer’s instructions. RNA integrity was corroborated as described before (Contreras et al., 2018). Two microgram RNA was reverse transcribed into cDNA using random primers and M-MLV reverse transcriptase (Invitrogen, CA, USA). RT-qPCR was performed in triplicate with the Eco Real-Time PCR System (Illumina, CA, USA), using primer sets for (Supplementary Table 1): *Tcf7, Lef1, Tcf7l1 (Tcf3), Tcf7l2 (Tcf4), Ccnd1* (*CyclinD1*), *Sox9, Axin2, Nfatc1, Ctnnb1* (β*-catenin*) and the housekeeping gene *18s* (used as a reference gene). The ΔΔCt method was used for quantification, and mRNA levels were expressed relative to the mean level of the control condition in each case. We analyzed and validated each RT-qPCR expected gene product using a 2% agarose gel. Digital droplet PCR was performed as previously described (Soliman et al., 2020). Gene expression analysis was performed using Taqman Gene Expression Assays (Applied Biosystems), on a 7900HT Real Time PCR system (Applied Biosystems). Sequence information for the primers contained in the Taqman assays is provided here: Taqman probes (Thermo Fisher Scientific): *Tcf7l2* mouse (Mm00501505_m1), *Tcf7l1* mouse (Mm01188711_m1), *Lef1* mouse (Mm00550265_m1), *Tcf7* mouse (Mm00493445_m1), *Runx2* mouse (Hs01047973_m1), *Adipoq* (Hs00605917_m1), and the housekeeping gene *Hprt* mouse (Mm03024075_m1). Data were acquired and analyzed using SDS 2.0 and SDS RQ Manager software (Applied Biosystems).

### Computational BioGRID database, Ubiquitination, and Tabula Muris open source database

The image in Fig. 6D was generated with BioGRID based on the human TCF7L2 interactome (Stark et al., 2006). The data from figures and tables in the BioGRID webpage (https://thebiogrid.org/) can be searched and sorted. For post-translational modification detection and delineation of ubiquitination at a site-specific level we used UbiSite webpage (http://csb.cse.yzu.edu.tw/UbiSite/) (Akimov et al., 2018). Murine Tabula Muris open database was used to generate the figures shown in Fig. S1E,F (The Tabula Muris Consortium et al., 2018). Data were extracted and analyzed from total tissues, diaphragm, and limb muscles.

### Statistical analysis

Mean ± s.e.m. values, as well as the number of experiments performed, are indicated in each figure. All datasets were analyzed for normal distribution using the D’agostino normality test. Statistical significance of the differences between the means was evaluated using the one-way analysis of variance (ANOVA) test followed by post-hoc Dunnett’s multiple comparison test and the significance level set at *P*<0.05. A two-tailed Student’s *t-* test was performed when two conditions were compared. Differences were considered significant with *P*<0.05. Data were collected in Microsoft Excel, and statistical analyses were performed using Prism 8 software for macOS (GraphPad).

## Abbreviations

CHX: Cycloheximide
CT: Connective tissue
DMD: Duchenne muscular dystrophy
ECM: Extracellular matrix
FAPs: Fibro-adipogenic progenitors
FACS: Fluorescence-activated cell sorting
HDACs: Histone deacetylases
IPF: Idiopathic pulmonary fibrosis
MSCs: Mesenchymal stromal cells
MI: Myocardial infarction
MuSCs: Muscle stem cells
MD: Muscular dystrophy
PDGFRα: Platelet-derived growth factor receptor alpha
Tcf/Lef: T-cell factor/lymphoid enhancer
TF: Transcription factor
TGF-β: Transforming growth factor type-beta
TSA: Trichostatin A
UPS: Ubiquitin-proteasome system

## Acknowledgments

We are grateful to the Unidad de Microscopía Avanzada (UMA) of Pontificia Universidad Católica de Chile for its support in image acquisition. We acknowledge the services provided by the UC CINBIOT Animal Facility funded by the PIA CONICYT* ECM-07 Program for Associative Research, of the Chilean National Council for Science and Technology. We also acknowledge the animal unit staff and genotyping core facility at the Biomedical Research Centre (UBC), especially Mr. Taka Murakami, Mrs. Krista Ranta, and Mr. Wei Yuan. We thank Mr. Andy Johnson and Mr. Justin Wong of UBC Flow Cytometry. We also acknowledge Martin Arostegui from Michael T. Underhill’s Lab (Biomedical Research Centre, UBC) for helping with MEFs culture. We acknowledge the Lab of Dr. Hugo Olguín (Pontificia Universidad Católica de Chile) for kindly providing USP7 small-molecule inhibitor HBX 41108 and also the Lab of Dr. Martín Montecino (Instituto de Ciencias Biomédicas, Universidad Andrés Bello) for kindly providing Trichostatin A. For administrative assistance, we thank Ms. Vanessa Morales and Mrs. Vittoria Canale. We finally acknowledge Mr. Eduardo Ramirez and Ms. Darling Vera for their technical support.

## Availability of data and materials

All data generated or analyzed during this study are included in this published article.

## Authors’ contributions

Conceptualization: O.C., E.B.; Methodology: O.C., H.S., M.T.; Software: O.C.; Validation: O.C., H.S., M.T., F.M.V.R., E.B.; Formal analysis: O.C., H.S.; Investigation: O.C., H.S., M.T.; Resources: E.B., F.M.V.R., O.C.; Data curation: O.C.; Writing - original draft: O.C.; Writing - review & editing: O.C., E.B., F.M.V.R., H.S., M.T.; Visualization: O.C., H.S., M.T., F.M.V.R., E.B.; Supervision: O.C., E.B., F.M.V.R.; Project administration: O.C., E.B.; Funding acquisition: E.B., F.M.V.R., O.C.

## Ethics approval and consent to participate

Not applicable.

## Consent for publication

Not applicable.

## References

Aberle, H., Bauer, A., Stappert, J., Kispert, A., & Kemler, R. (1997). β-catenin is a target for the ubiquitin– proteasome pathway. The EMBO Journal, 16(13), 3797–3804. https://doi.org/10.1093/emboj/16.13.3797

Accornero, F., O. Kanisicak, A. Tjondrokoesoemo, A.C. Attia, E.M. McNally, and J.D. Molkentin. 2014. Myofiber-specific inhibition of TGFbeta signaling protects skeletal muscle from injury and dystrophic disease in mice. Hum Mol Genet. 23:6903–6915.

Bettica, P., Petrini, S., D’Oria, V., D’Amico, A., Catteruccia, M., Pane, M., … Mercuri, E. (2016). Histological effects of givinostat in boys with Duchenne muscular dystrophy. Neuromuscular Disorders, 26(10), 643–649. https://doi.org/10.1016/j.nmd.2016.07.002

Acuña, M.J., P. Pessina, H. Olguin, D. Cabrera, C.P. Vio, M. Bader, P. Munoz-Canoves, R.A. Santos, C. Cabello-Verrugio, and E. Brandan. 2014. Restoration of muscle strength in dystrophic muscle by angiotensin-1-7 through inhibition of TGF-beta signalling. Hum Mol Genet. 23:1237–1249.

Agley, C. C., Rowlerson, A. M., Velloso, C. P., Lazarus, N. R., & Harridge, S. D. R. (2013). Human skeletal muscle fibroblasts, but not myogenic cells, readily undergo adipogenic differentiation. Journal of Cell Science, 126(24), 5610 LP – 5625. https://doi.org/10.1242/jcs.132563

Akimov, V., Barrio-Hernandez, I., Hansen, S. V. F., Hallenborg, P., Pedersen, A.-K., Bekker-Jensen, D. B., … Blagoev, B. (2018). UbiSite approach for comprehensive mapping of lysine and N-terminal ubiquitination sites. Nature Structural & Molecular Biology, 25(7), 631–640. https://doi.org/10.1038/s41594-018-0084-y

Asli, N. S., Xaymardan, M., Patrick, R., Farbehi, N., Cornwell, J., Forte, E., … Harvey, R. P. (2019). PDGFRα signaling in cardiac fibroblasts modulates quiescence, metabolism and self-renewal, and promotes anatomical and functional repair. BioRxiv, 225979. https://doi.org/10.1101/225979

Burgy, O., & Königshoff, M. (2018). The WNT signaling pathways in wound healing and fibrosis. Matrix biol, 68–69, 67–80. https://doi.org/10.1016/j.matbio.2018.03.017

Bernasconi, P., C. Di Blasi, M. Mora, L. Morandi, S. Galbiati, P. Confalonieri, F. Cornelio, and R. Mantegazza. 1999. Transforming growth factor-beta1 and fibrosis in congenital muscular dystrophies. Neuromuscul Disord. 9:28–33.

Biressi, S., Miyabara, E. H., Gopinath, S. D., Carlig, P. M. M., & Rando, T. A. (2014). A Wnt-TGF2 axis induces a fibrogenic program in muscle stem cells from dystrophic mice. Science Translational Medicine. https://doi.org/10.1126/scitranslmed.3008411

Bolden, J. E., Peart, M. J., & Johnstone, R. W. (2006). Anticancer activities of histone deacetylase inhibitors. Nature Reviews Drug Discovery, 5(9), 769–784. https://doi.org/10.1038/nrd2133

Brack, A. S., Conboy, M. J., Roy, S., Lee, M., Kuo, C. J., Keller, C., & Rando, T. A. (2007). Increased Wnt signaling during aging alters muscle stem cell fate and increases fibrosis. Science. https://doi.org/10.1126/science.1144090

Braun, T., G. Buschhausen-Denker, E. Bober, E. Tannich, and H.H. Arnold. 1989. A novel human muscle factor related to but distinct from MyoD1 induces myogenic conversion in 10T1/2 fibroblasts. EMBO J. 8:701–709.

Callahan, J.F., J.L. Burgess, J.A. Fornwald, L.M. Gaster, J.D. Harling, F.P. Harrington, J. Heer, C. Kwon, R. Lehr, A. Mathur, B.A. Olson, J. Weinstock, and N.J. Laping. 2002. Identification of novel inhibitors of the transforming growth factor beta1 (TGF-beta1) type 1 receptor (ALK5). J Med Chem. 45:999–1001.

Cao, H., Wang, C., Chen, X., Hou, J., Xiang, Z., Shen, Y., & Han, X. (2018). Inhibition of Wnt/β-catenin signaling suppresses myofibroblast differentiation of lung resident mesenchymal stem cells and pulmonary fibrosis. Scientific Reports, 8(1), 13644. https://doi.org/10.1038/s41598-018-28968-9

Carr, M. J., Toma, J. S., Johnston, A. P. W., Steadman, P. E., Yuzwa, S. A., Mahmud, N., … Miller, F. D. (2019). Mesenchymal Precursor Cells in Adult Nerves Contribute to Mammalian Tissue Repair and Regeneration. Cell Stem Cell, 24(2), 240-256.e9. https://doi.org/10.1016/j.stem.2018.10.024

Ceco, E., and E.M. McNally. 2013. Modifying muscular dystrophy through transforming growth factor-beta. FEBS J. 280:4198–4209.

Chen, X., Ayala, I., Shannon, C., Fourcaudot, M., Acharya, N. K., Jenkinson, C. P., … Norton, L. (2018). The Diabetes Gene and Wnt Pathway Effector TCF7L2 Regulates Adipocyte Development and Function. Diabetes, 67(4), 554 LP – 568. https://doi.org/10.2337/db17-0318

Chong J.J.H. Chandrakanthan V. Xaymardan M. Asli N.S. Li J. Ahmed I. Heffernan C. Menon M.K. Scarlett C.J. Rashidianfar A. et al. Adult cardiac-resident MSC-like stem cells with a proepicardial origin. Cell Stem Cell. 2011; 9: 527–540

Chilosi, M. et al. Aberrant Wnt/beta-catenin pathway activation in idiopathic pulmonary fibrosis. Am. J. Pathol. 162, 1495–1502 (2003).

Cho, N., Razipour, S. E., & McCain, M. L. (2018). TGF-β1 dominates extracellular matrix rigidity for inducing differentiation of human cardiac fibroblasts to myofibroblasts. Experimental Biology and Medicine, 243(7), 601–612. https://doi.org/10.1177/1535370218761628

Cisternas, P., Henriquez, J. P., Brandan, E., & Inestrosa, N. C. (2014). Wnt Signaling in Skeletal Muscle Dynamics: Myogenesis, Neuromuscular Synapse and Fibrosis. Molecular Neurobiology, 49(1), 574–589. https://doi.org/10.1007/s12035-013-8540-5

Clevers, H., 2006. Wnt/β-Catenin Signaling in Development and Disease. Cell 127: 3, 469-480.

Cohn, R.D., C. van Erp, J.P. Habashi, A.A. Soleimani, E.C. Klein, M.T. Lisi, M. Gamradt, C.M. ap Rhys, T.M. Colland, F., Formstecher, E., Jacq, X., Reverdy, C., Planquette, C., Conrath, S., … Daviet, L. (2009). Small-molecule inhibitor of USP7/HAUSP ubiquitin protease stabilizes and activates p53 in cells. Molecular Cancer Therapeutics, 8(8), 2286 LP – 2295. https://doi.org/10.1158/1535-7163.MCT-09-0097

Holm, B.L. Loeys, F. Ramirez, D.P. Judge, C.W. Ward, and H.C. Dietz. 2007. Angiotensin II type 1 receptor blockade attenuates TGF-beta-induced failure of muscle regeneration in multiple myopathic states. Nat Med. 13:204–210.

Colwell, A. S., Krummel, T. M., Longaker, M. T. & Lorenz, H. P. Wnt-4 expression is increased in fibroblasts after TGF-beta1 stimulation and during fetal and postnatal wound repair. Plast. Reconstr. Surg. 117, 2297–2301 (2006).

Contreras, O., D.L. Rebolledo, J.E. Oyarzun, H.C. Olguin, and E. Brandan. 2016. Connective tissue cells expressing fibro/adipogenic progenitor markers increase under chronic damage: relevance in fibroblast-myofibroblast differentiation and skeletal muscle fibrosis. Cell Tissue Res. 364:647–660.

Contreras, O., M. Villarreal, and E. Brandan. 2018. Nilotinib impairs skeletal myogenesis by increasing myoblast proliferation. Skelet Muscle 8, 5. doi: 10.1186/s13395-018-0150-5

Contreras O, Rebolledo D, Oyarzún J and Brandan E. Fibroblasts (Tcf4) and mesenchymal progenitors (PDGFRα) correspond to the same cell type and are increased in skeletal muscle dystrophy, denervation and chronic damage [version 1; not peer reviewed]. F1000Research 2019a, 8:299 (poster) (https://doi.org/10.7490/f1000research.1116478.1)

Contreras, O., Rossi, F. M., & Brandan, E. (2019b). Adherent muscle connective tissue fibroblasts are phenotypically and biochemically equivalent to stromal fibro/adipogenic progenitors. Matrix Biology Plus. https://doi.org/10.1016/j.mbplus.2019.04.003

Contreras, O., Cruz-Soca, M., Theret, M., Soliman, H., Tung, L. W., Groppa, E., Rossi, F. M. and Brandan, E. (2019c). Cross-talk between TGF-β and PDGFRα signaling pathways regulates the fate of stromal fibro– adipogenic progenitors. Journal of Cell Science. 132, 232157. https://doi.org/10.1242/JCS.232157

Contreras, O. (2019d). Hic1 deletion unleashes quiescent connective tissue stem cells and impairs skeletal muscle regeneration. Journal of Cell Communication and Signaling. https://doi.org/10.1007/s12079-019-00545-3

Cosin-Roger, J., Ortiz-Masià, M. D., & Barrachina, M. D. (2019). Macrophages as an Emerging Source of Wnt Ligands: Relevance in Mucosal Integrity. Frontiers in Immunology. Retrieved from https://www.frontiersin.org/article/10.3389/fimmu.2019.02297

Danna, N.R., B.G. Beutel, K.A. Campbell, and J.A. Bosco, 3rd. 2014. Therapeutic approaches to skeletal muscle repair and healing. Sports Health. 6:348–355.

David, C. J. and Massagué, J. (2018). Contextual determinants of TGFβ action in development, immunity and cancer. Nat. Rev. Mol. Cell Biol. 19, 419–435. doi: 10.1038/s41580-018-0007-0

de la Vega, E., González, N., Cabezas, F., Montecino, F., Blanco, N., & Olguín, H. (2020). Usp7-Dependent control of myogenin stability is required for terminal differentiation in skeletal muscle progenitors. The FEBS Journal. Accepted Author Manuscript. https://doi.org/10.1111/febs.15269

Derynck, R., & Budi, E. H. (2019). Specificity, versatility, and control of TGF-b family signaling. Science Signaling. https://doi.org/10.1126/scisignal.aav5183

Driskell, R. R., Lichtenberger, B. M., Hoste, E., Kretzschmar, K., Simons, B. D., Charalambous, M., Ferron, S. R., Herault, Y., Pavlovic, G., Ferguson-Smith, A. C. et al. (2013). Distinct fibroblast lineages determine dermal architecture in skin development and repair. Nature 504, 277–281. doi: 10.1038/nature12783

Droguett, R., Cabello-Verrugio, C., Riquelme, C., & Brandan, E. (2006). Extracellular proteoglycans modify TGF-β bio-availability attenuating its signaling during skeletal muscle differentiation. Matrix biol. https://doi.org/10.1016/j.matbio.2006.04.004

Dulauroy S, Di Carlo SE, Langa F, Eberl G, Peduto L. Lineage tracing and genetic ablation of ADAM12(+) perivascular cells identify a major source of profibrotic cells during acute tissue injury. Nature medicine. 2012;18(8):1262–70. pmid:22842476

Dzialo, E., Tkacz, K., & Błyszczuk, P. (2018). Crosstalk between the TGF-β and WNT signalling pathways during cardiac fibrogenesis. Acta Biochimica Polonica. https://doi.org/10.18388/abp.2018_2635

Elfert, S., Weise, A., Bruser, K., Biniossek, M. L., Jägle, S., Senghaas, N., & Hecht, A. (2013). Acetylation of Human TCF4 (TCF7L2) Proteins Attenuates Inhibition by the HBP1 Repressor and Induces a Conformational Change in the TCF4::DNA Complex. PLoS ONE. https://doi.org/10.1371/journal.pone.0061867

Farbehi, N., Patrick, R., Dorison, A., Xaymardan, M., Janbandhu, V., Wystub-Lis, K., Ho, J. W. K., Nordon, R. E. and Harvey, R. P. (2019). Single-cell expression profiling reveals dynamic flux of cardiac stromal, vascular and immune cells in health and injury. eLife 8, e43882. doi: 10.7554/eLife.43882

Frietze, S., Wang, R., Yao, L., Tak, Y., Ye, Z., Gaddis, M., Witt, H., Farnham, P., and Jin, V., 2012. Cell type-specific binding patterns reveal that TCF7L2 can be tethered to the genome by association with GATA3. Genome Biology 13, R52.

Fu, X., Khalil, H., Kanisicak, O., Boyer, J. G., Vagnozzi, R. J., Maliken, B. D., … Molkentin, J. D. (2018). Specialized fibroblast differentiated states underlie scar formation in the infarcted mouse heart. The Journal of Clinical Investigation, 128(5), 2127–2143. https://doi.org/10.1172/JCI98215

Furtado, M. B., Nim, H. T., Boyd, S. E. and Rosenthal, N. A. (2016). View from the heart: cardiac fibroblasts in development, scarring and regeneration. Development 143, 387–397. doi: 10.1242/dev.120576

Girardi, F., & Le Grand, F. (2018). Chapter Five - Wnt Signaling in Skeletal Muscle Development and Regeneration. In J. Larraín & G. B. T.-P. in M. B. and T. S. Olivares (Eds.), WNT Signaling in Health and Disease (Vol. 153, pp. 157–179). Academic Press. https://doi.org/10.1016/bs.pmbts.2017.11.026

Gonzalez, D., O. Contreras, D.L. Rebolledo, J.P. Espinoza, B. van Zundert, and E. Brandan. 2017. ALS skeletal muscle shows enhanced TGF-beta signaling, fibrosis and induction of fibro/adipogenic progenitor markers. PLoS One. 12: e0177649.

Gosselin, L. E., Williams, J. E., Deering, M., Brazeau, D., Koury, S., & Martinez, D. A. (2004). Localization and early time course of TGF-β1 mRNA expression in dystrophic muscle. Muscle and Nerve. https://doi.org/10.1002/mus.20150

Götze, S., Coersmeyer, M., Müller, O., & Sievers, S. (2014). Histone deacetylase inhibitors induce attenuation of Wnt signaling and TCF7L2 depletion in colorectal carcinoma cells. International Journal of Oncology, 45, 1715–1723. https://doi.org/10.3892/ijo.2014.2550

Grant, S. F. A., Thorleifsson, G., Reynisdottir, I., Benediktsson, R., Manolescu, A., Sainz, J., … Stefansson, K. (2006). Variant of transcription factor 7-like 2 (TCF7L2) gene confers risk of type 2 diabetes. Nature Genetics, 38, 320–323. https://doi.org/10.1038/ng1732

Greer, C. B., Tanaka, Y., Kim, Y. J., Xie, P., Zhang, M. Q., Park, I.-H., & Kim, T. H. (2015). Histone Deacetylases Positively Regulate Transcription through the Elongation Machinery. Cell Reports, 13(7), 1444–1455. https://doi.org/10.1016/j.celrep.2015.10.013

Gutierrez, J., C.A. Droppelmann, O. Contreras, C. Takahashi, and E. Brandan. 2015. RECK-Mediated beta1-Integrin Regulation by TGF-beta1 Is Critical for Wound Contraction in Mice. PLoS One. 10: e0135005.

Hamburg-Shields, E., DiNuoscio, G. J., Mullin, N. K., Lafyatis, R., & Atit, R. P. (2015). Sustained β-catenin activity in dermal fibroblasts promotes fibrosis by up-regulating expression of extracellular matrix protein-coding genes. The Journal of Pathology, 235(5), 686–697. https://doi.org/10.1002/path.4481

Hamilton, T.G., R.A. Klinghoffer, P.D. Corrin, and P. Soriano. 2003. Evolutionary divergence of platelet-derived growth factor alpha receptor signaling mechanisms. Mol Cell Biol. 23: 4013–4025.

He, W., Zhang, L., Ni, A., Zhang, Z., Mirotsou, M., Mao, L., … Dzau, V. J. (2010). Exogenously administered secreted frizzled related protein 2 (Sfrp2) reduces fibrosis and improves cardiac function in a rat model of myocardial infarction. Proceedings of the National Academy of Sciences, 107(49), 21110 LP – 21115. https://doi.org/10.1073/pnas.1004708107

He, W., Dai, C., Li, Y., Zeng, G., Monga, S. P., & Liu, Y. (2009). Wnt/β-Catenin Signaling Promotes Renal Interstitial Fibrosis. Journal of the American Society of Nephrology, 20(4), 765 LP – 776. https://doi.org/10.1681/ASN.2008060566

Henderson, W. R., Chi, E. Y., Ye, X., Nguyen, C., Tien, Y., Zhou, B., … Kahn, M. (2010). Inhibition of Wnt/β-catenin/CREB binding protein (CBP) signaling reverses pulmonary fibrosis. Proceedings of the National Academy of Sciences, 107(32), 14309 LP – 14314. https://doi.org/10.1073/pnas.1001520107

Heredia, J.E., L. Mukundan, F.M. Chen, A.A. Mueller, R.C. Deo, R.M. Locksley, T.A. Rando, and A. Chawla. 2013. Type 2 innate signals stimulate fibro/adipogenic progenitors to facilitate muscle regeneration. Cell. 153:376–388.

Ieronimakis, N., Hays, A., Prasad, A., Janebodin, K., Duffield, J. S. and Reyes, M. (2016). PDGFRα signalling promotes fibrogenic responses in collagenproducing cells in Duchenne muscular dystrophy. J. Pathol. 240, 410–424. doi: 10.1002/path.4801

Ishitani, T., Ninomiya-Tsuji, J., & Matsumoto, K. (2003). Regulation of Lymphoid Enhancer Factor 1/T-Cell Factor by Mitogen-Activated Protein Kinase-Related Nemo-Like Kinase-Dependent Phosphorylation in Wnt/β-Catenin Signaling. Molecular and Cellular Biology, 23(4), 1379 LP–1389. https://doi.org/10.1128/MCB.23.4.1379-1389.2003

Jin, T. (2016). Current Understanding on Role of the Wnt Signaling Pathway Effector TCF7L2 in Glucose Homeostasis. Endocrine Reviews, 37(3), 254–277. https://doi.org/10.1210/er.2015-1146

Joe, A.W., L. Yi, A. Natarajan, F. Le Grand, L. So, J. Wang, M.A. Rudnicki, and F.M. Rossi. 2010. Muscle injury activates resident fibro/adipogenic progenitors that facilitate myogenesis. Nat Cell Biol. 12:153–163.

Jones, D. L., Haak, A. J., Caporarello, N., Choi, K. M., Ye, Z., Yan, H., … Tschumperlin, D. J. (2019). TGFβ-induced fibroblast activation requires persistent and targeted HDAC-mediated gene repression. Journal of Cell Science, 132(20), jcs233486. https://doi.org/10.1242/jcs.233486

Judson, R. N., Low, M., Eisner, C., & Rossi, F. M. (2017). Isolation, Culture, and Differentiation of Fibro/Adipogenic Progenitors (FAPs) from Skeletal Muscle. In Methods in molecular biology (Clifton, N.J.). https://doi.org/10.1007/978-1-4939-7283-8_7

Kanisicak, O., Khalil, H., Ivey, M. J., Karch, J., Maliken, B. D., Correll, R. N., Brody, M. J., J. Lin, S.-C., Aronow, B. J., Tallquist, M. D. et al. (2016). Genetic lineage tracing defines myofibroblast origin and function in the injured heart. Nat. Commun. 7, 12260. doi: 10.1038/ncomms12260

Kardon, G., Harfe, B. D., & Tabin, C. J. (2003). A Tcf4-Positive Mesodermal Population Provides a Prepattern for Vertebrate Limb Muscle Patterning. Developmental Cell, 5(6), 937–944. https://doi.org/10.1016/S1534-5807(03)00360-5

Kim, K.K., D. Sheppard, and H.A. Chapman. 2018. TGF-beta1 Signaling and Tissue Fibrosis. Cold Spring Harb Perspect Biol. 10.

Konigsho, M. et al. Functional Wnt signaling is increased in idiopathic pulmonary fibrosis. PLoS One. 3, e2142 (2008).

Kopinke, D., E.C. Roberson, and J.F. Reiter. 2017. Ciliary Hedgehog Signaling Restricts Injury-Induced Adipogenesis. Cell. 170:340–351 e312.

Korinek V, Barker N, Moerer P & van Donselaar E., 1998. Depletion of epithelial stem-cell compartments in the small intestine of mice lacking Tcf-4. Nature genetics. 18, 379–383.

Korinek, V, Barker, N., Morin, P.J., van Wichen, D., de Weger, R., Kinzler, K.W., Vogelstein, B., and Clevers, H., 1997. Constitutive Transcriptional Activation by a beta-catenin-Tcf complex in APC -/- Colon Carcinoma. Science, 275: 1784–1787

Lemos, D.R., F. Babaeijandaghi, M. Low, C.K. Chang, S.T. Lee, D. Fiore, R.H. Zhang, A. Natarajan, S.A. Nedospasov, and F.M. Rossi. 2015. Nilotinib reduces muscle fibrosis in chronic muscle injury by promoting TNF-mediated apoptosis of fibro/adipogenic progenitors. Nat Med. 21:786–794.

Lemos, D. R. and Duffield, J. S. (2018). Tissue-resident mesenchymal stromal cells: Implications for tissue-specific antifibrotic therapies. Science Translational Medicine. doi: 10.1126/scitranslmed.aan5174

Lepper, C., T.A. Partridge, and C.M. Fan. 2011. An absolute requirement for Pax7-positive satellite cells in acute injury-induced skeletal muscle regeneration. Development. 138:3639–3646.

Lien WH, Fuchs E., 2014. Wnt some lose some: transcriptional governance of stem cells by Wnt/beta-catenin signaling. Genes Dev. 28:1517–1532.

Lien, W. H., Polak, L., Lin, M., Lay, K., Zheng, D., & Fuchs, E. (2014). In vivo transcriptional governance of hair follicle stem cells by canonical Wnt regulators. Nature Cell Biology. https://doi.org/10.1038/ncb2903

Liu, L. et al. Wnt pathway in pulmonary fibrosis in the bleomycin mouse model. J. Environ. Pathol. Toxicol. Oncol. 28, 99–108 (2009).

Lukjanenko, L., Karaz, S., Stuelsatz, P., Rudnicki, M. A., Bentzinger, C. F., Feige, J. N., … Michaud, J. (2019). Aging Disrupts Muscle Stem Cell Function by Impairing Matricellular WISP1 Secretion from Fibro-Adipogenic Progenitors Stem Cell, 1–14. https://doi.org/10.1016/j.stem.2018.12.014

Lynch, M. D. and Watt, F. M. (2018). Fibroblast heterogeneity: implications for human disease. J. Clin. Investig. 128, 26–35. doi: 10.1172/JCI93555

Mackey, A.L., M. Magnan, B. Chazaud, and M. Kjaer. 2017. Human skeletal muscle fibroblasts stimulate in vitro myogenesis and in vivo muscle regeneration. J Physiol. 595:5115–5127.

Malecova, B., S. Gatto, U. Etxaniz, M. Passafaro, A. Cortez, C. Nicoletti, L. Giordani, A. Torcinaro, M. De Bardi, S. Bicciato, F. De Santa, L. Madaro, and P.L. Puri. 2018. Dynamics of cellular states of fibro-adipogenic progenitors during myogenesis and muscular dystrophy. Nat Commun. 9:3670.

Mahmoudi, S., Mancini, E., Xu, L., Moore, A., Jahanbani, F., Hebestreit, K., … Brunet, A. (2019). Heterogeneity in old fibroblasts is linked to variability in reprogramming and wound healing. Nature, 574(7779), 553–558. https://doi.org/10.1038/s41586-019-1658-5

Mann, C.J., E. Perdiguero, Y. Kharraz, S. Aguilar, P. Pessina, A.L. Serrano, and P. Munoz-Canoves. 2011. Aberrant repair and fibrosis development in skeletal muscle. Skelet Muscle. 1:21.

Masur, S., & Dewal, H. (1996). Myofibroblasts differentiate from fibroblasts when plated at low density. Proceedings of the National Academy of Sciences of the United States of America. https://doi.org/10.1073/pnas.93.9.4219

Massagué, J., Cheifetz, S., Endo, T., & Nadal-Ginard, B. (1986). Type beta transforming growth factor is an inhibitor of myogenic differentiation. Proceedings of the National Academy of Sciences, 83(21), 8206 LP–8210. https://doi.org/10.1073/pnas.83.21.8206

Massagué, J. (1998). TGF-beta signal transduction. Annu Rev Biochem. https://doi.org/10.1146/annurev.biochem.67.1.753

Massagué, J. (2012). TGFβ signalling in context. Nature Reviews Molecular Cell Biology. https://doi.org/10.1038/nrm3434

Mathew, S.J., J.M. Hansen, A.J. Merrell, M.M. Murphy, J.A. Lawson, D.A. Hutcheson, M.S. Hansen, M. Angus-Hill, and G. Kardon. 2011. Connective tissue fibroblasts and Tcf4 regulate myogenesis. Development. 138: 371–384.

McBeath, R., Pirone, D. M., Nelson, C. M., Bhadriraju, K., & Chen, C. S. (2004). Cell shape, cytoskeletal tension, and RhoA regulate stem cell lineage commitment. Developmental Cell. https://doi.org/10.1016/S1534-5807(04)00075-9

Merrell, A. J., Ellis, B. J., Fox, Z. D., Lawson, J. A., Weiss, J. A., & Kardon, G. (2015). Muscle connective tissue controls development of the diaphragm and is a source of congenital diaphragmatic hernias. Nature Genetics. https://doi.org/10.1038/ng.3250

Minetti, G., Colussi, C., Adami, R. et al. Functional and morphological recovery of dystrophic muscles in mice treated with deacetylase inhibitors. Nat Med 12, 1147–1150 (2006). https://doi.org/10.1038/nm1479

Mozzetta, C., Consalvi, S., Saccone, V., Tierney, M., Diamantini, A., Mitchell, K.J., Marazzi, G., Borsellino, G., Battistini, L., Sassoon, D., Sacco, A. and Puri, P.L. (2013), Fibroadipogenic progenitors mediate the ability of HDAC inhibitors to promote regeneration in dystrophic muscles of young, but not old Mdx mice. EMBO Mol Med, 5: 626–639. doi: 10.1002/emmm.201202096

Murphy, M.M., J.A. Lawson, S.J. Mathew, D.A. Hutcheson, and G. Kardon. 2011. Satellite cells, connective tissue fibroblasts and their interactions are crucial for muscle regeneration. Development. 138:3625–3637.

Nakano, N., Itoh, S., Watanabe, Y., Maeyama, K., Itoh, F., & Kato, M. (2010). Requirement of TCF7L2 for TGF-β-dependent transcriptional activation of the TMEPAI gene. Journal of Biological Chemistry. https://doi.org/10.1074/jbc.M110.132209

Nalepa, G., Rolfe, M., & Harper, J. W. (2006). Drug discovery in the ubiquitin-proteasome system. Nature Reviews Drug Discovery. https://doi.org/10.1038/nrd2056

Nawshad, A., Medici, D., Liu, C.-C., & Hay, E. D. (2007). TGFβ3 inhibits E-cadherin gene expression in palate medial-edge epithelial cells through a Smad2-Smad4-LEF1 transcription complex. Journal of Cell Science, 120(9), 1646 LP – 1653. https://doi.org/10.1242/jcs.003129

Nusse, R., & Clevers, H. (2017). Wnt/β-Catenin Signaling, Disease, and Emerging Therapeutic Modalities. Cell. https://doi.org/10.1016/j.cell.2017.05.016

Oishi, T., Uezumi, A., Kanaji, A., Yamamoto, N., Yamaguchi, A., Yamada, H., et al. (2013). Osteogenic differentiation capacity of human skeletal muscle-derived progenitor cells. PLoS ONE 8:e56641. doi: 10.1371/journal.pone.0056641

Pessina, P., Y. Kharraz, M. Jardi, S. Fukada, A.L. Serrano, E. Perdiguero, and P. Munoz-Canoves. 2015. Fibrogenic Cell Plasticity Blunts Tissue Regeneration and Aggravates Muscular Dystrophy. Stem Cell Reports. 4:1046–1060.

Piersma, B., Bank, R. A., & Boersema, M. (2015). Signaling in Fibrosis: TGF-β, WNT, and YAP/TAZ Converge. Frontiers in Medicine. https://doi.org/10.3389/fmed.2015.00059

Plikus, M. V., Guerrero-Juarez, C. F., Ito, M., Li, Y. R., Dedhia, P. H., Zheng, Y., Shao, M., Gay, D. L., Ramos, R., Hsi, T.-C. et al. (2017). Regeneration of fat cells from myofibroblasts during wound healing. Science 355, 748–752. doi: 10.1126/science.aai8792

Ravindranath A, O’Connell A, Johnston PG, El-Tanani MK., 2008. The role of LEF/ TCF factors in neoplastic transformation. Curr. Mol. Med. 8, 38–50.

Reznikoff, C.A., J.S. Bertram, D.W. Brankow, and C. Heidelberger. 1973. Quantitative and qualitative studies of chemical transformation of cloned C3H mouse embryo cells sensitive to post-confluence inhibition of cell division. Cancer Res. 33:3239–3249.

Rinkevich, Y.,Walmsley, G. G., Hu, M. S., Maan, Z. N., Newman, A. M., Drukker, M., Januszyk, M., Krampitz, G. W., Gurtner, G. C., Lorenz, H. P. et al. (2015). Identification and isolation of a dermal lineage with intrinsic fibrogenic potential. Science 348, aaa2151. doi: 10.1126/science.aaa2151

Riquelme, C., Larraín, J., Schönherr, E., Henriquez, J. P., Kresse, H., & Brandan, E. (2001). Antisense Inhibition of Decorin Expression in Myoblasts Decreases Cell Responsiveness to Transforming Growth Factor α and Accelerates Skeletal Muscle Differentiation. Journal of Biological Chemistry. https://doi.org/10.1074/jbc.M004602200

Riquelme-Guzmán, C., O. Contreras, and E. Brandan. 2018. Expression of CTGF/CCN2 in response to LPA is stimulated by fibrotic extracellular matrix via the integrin/FAK axis. Am J Physiol Cell Physiol. 314:C415–C427.

Riquelme-Guzmán, C., & Contreras, O. (2020). Single-cell revolution unveils the mysteries of the regenerative mammalian digit tip. Developmental Biology. https://doi.org/10.1016/j.ydbio.2020.02.002

Rognoni, E., Pisco, A. O., Hiratsuka, T., SipilalJ, K. H., Belmonte, J. M., Mobasseri, S. A., Philippeos, C., Dilão, R. and Watt, F. M. (2018). Fibroblast state switching orchestrates dermal maturation and wound healing. Mol. Syst. Biol. 14, e8174. doi: 10.15252/msb.20178174

Saccone, V., Consalvi, S., Giordani, L., Mozzetta, C., Barozzi, I., Sandoná, M., … Puri, P. L. (2014). HDAC-regulated myomiRs control BAF60 variant exchange and direct the functional phenotype of fibro-adipogenic progenitors in dystrophic muscles. Genes & Development, 28(8), 841–857. https://doi.org/10.1101/gad.234468.113

Sambasivan, R., R. Yao, A. Kissenpfennig, L. Van Wittenberghe, A. Paldi, B. Gayraud-Morel, H. Guenou, B. Malissen, S. Tajbakhsh, and A. Galy. 2011. Pax7-expressing satellite cells are indispensable for adult skeletal muscle regeneration. Development. 138:3647–3656.

Schabort, E. J., van der Merwe, M., Loos, B., Moore, F. P., & Niesler, C. U. (2009). TGF-β’s delay skeletal muscle progenitor cell differentiation in an isoform-independent manner. Experimental Cell Research, 315(3), 373–384. https://doi.org/10.1016/j.yexcr.2008.10.037

Schuijers, J., Mokry, M., Hatzis, P., Cuppen, E., and Clevers, H., 2014. Wnt-induced transcriptional activation is exclusively mediated by TCF/LEF. EMBO J, 33, 146–156.

Scott, R. W., Arostegui, M., Schweitzer, R., Rossi, F. M. V., & Underhill, T. M. (2019). Hic1 Defines Quiescent Mesenchymal Progenitor Subpopulations with Distinct Functions and Fates in Skeletal Muscle Regeneration. Cell Stem Cell. https://doi.org/10.1016/j.stem.2019.11.004

Seo, H.-H., Lee, S., Lee, C. Y., Lee, J., Shin, S., Song, B.-W., … Hwang, K.-C. (2019). Multipoint targeting of TGF-β/Wnt transactivation circuit with microRNA 384-5p for cardiac fibrosis. Cell Death & Differentiation, 26(6), 1107–1123. https://doi.org/10.1038/s41418-018-0187-3

Seto, E., & Yoshida, M. (2014). Erasers of Histone Acetylation: The Histone Deacetylase Enzymes. Cold Spring Harbor Perspectives in Biology, 6(4). https://doi.org/10.1101/cshperspect.a018713

Schafer, S., Viswanathan, S., Widjaja, A. et al. IL-11 is a crucial determinant of cardiovascular fibrosis. Nature 552, 110–115 (2017). https://doi.org/10.1038/nature24676

Shy, B. R., Wu, C. I., Khramtsova, G. F., Zhang, J. Y., Olopade, O. I., Goss, K. H., & Merrill, B. J. (2013). Regulation of Tcf7l1 DNA Binding and Protein Stability as Principal Mechanisms of Wnt/β-Catenin Signaling. Cell Reports. https://doi.org/10.1016/j.celrep.2013.06.001

Singh, R., J.N. Artaza, W.E. Taylor, N.F. Gonzalez-Cadavid, and S. Bhasin. 2003. Androgens stimulate myogenic differentiation and inhibit adipogenesis in C3H/10T1/2 pluripotent cells through an androgen receptor-mediated pathway. Endocrinology. 144:5081–5088.

Smith, L. R. and Barton, E. R. (2018). Regulation of fibrosis in muscular dystrophy. Matrix Biol. 68-69, 602–615. doi: 10.1016/j.matbio.2018.01.014

Song, Y., Lee, S., Kim, J.R., Jho, E-H. (2018). Pja2 Inhibits Wnt/β-catenin Signaling by Reducing the Level of TCF/LEF1. International Journal of Stem Cells, 11(2), 242–247. https://doi.org/10.15283/ijsc18032

Soliman, H., Paylor, B., Scott, R. W., Lemos, D. R., Chang, C., Arostegui, M., … Rossi, F. M. V. (2020). Pathogenic Potential of Hic1-Expressing Cardiac Stromal Progenitors. Cell Stem Cell, 26(2), 205-220.e8. https://doi.org/10.1016/j.stem.2019.12.008

Stark, C. (2006). BioGRID: a general repository for interaction datasets. Nucleic Acids Research. https://doi.org/10.1093/nar/gkj109

Surendran, K., McCaul, S. P. & Simon, T. C. A role for Wnt-4 in renal fibrosis. Am. J. Physiol. Renal. Physiol. 282, F431–F441 (2002).

Suzuki, K., Bose, P., Leong-Quong, R. Y., Fujita, D. J., & Riabowol, K. (2010). REAP: A two minute cell fractionation method. BMC Research Notes. https://doi.org/10.1186/1756-0500-3-294

Tang, W., Dodge, M., Gundapaneni, D., Michnoff, C., Roth, M., & Lum, L. (2008). A genome-wide RNAi screen for Wnt/β-catenin pathway components identifies unexpected roles for TCF transcription factors in cancer. Proceedings of the National Academy of Sciences, 105(28), 9697 LP – 9702. https://doi.org/10.1073/pnas.0804709105

Tallquist, M. D., & Molkentin, J. D. (2017). Redefining the identity of cardiac fibroblasts. Nature Reviews Cardiology. https://doi.org/10.1038/nrcardio.2017.57

The Tabula Muris Consortium, Overall coordination, Logistical coordination, Organ collection and processing; Library preparation and sequencing, Computational data analysis, Cell type annotation, Writing group, Supplemental text writing group and Principal investigators. (2018). Single-cell transcriptomics of 20 mouse organs creates a Tabula Muris. Nature 562, 367–372. doi: 10.1038/s41586-018-0590-4

Tian, X., Zhang, J., Tan, T. K., Lyons, J. G., Zhao, H., Niu, B., … Harris, D. C. H. (2013). Association of β-catenin with P-Smad3 but not LEF-1 dissociates in vitro profibrotic from anti-inflammatory effects of TGF-β1. Journal of Cell Science, 126(1), 67–76. https://doi.org/10.1242/jcs.103036

Trensz, F., Haroun, S., Cloutier, A., Richter, M. V. & Grenier, G. A muscle resident cell population promotes fibrosis in hindlimb skeletal muscles of mdx mice through the Wnt canonical pathway. Am. J. Physiol. Cell Physiol. 299, C939–C947 (2010).

Uezumi, A., S. Fukada, N. Yamamoto, M. Ikemoto-Uezumi, M. Nakatani, M. Morita, A. Yamaguchi, H. Yamada, I. Nishino, Y. Hamada, and K. Tsuchida. 2014a. Identification and characterization of PDGFRalpha+ mesenchymal progenitors in human skeletal muscle. Cell Death Dis. 5: e1186.

Uezumi, A., M. Ikemoto-Uezumi, and K. Tsuchida. 2014b. Roles of nonmyogenic mesenchymal progenitors in pathogenesis and regeneration of skeletal muscle. Front Physiol. 5:68.

Uezumi, A., S. Fukada, N. Yamamoto, S. Takeda, and K. Tsuchida. 2010. Mesenchymal progenitors distinct from satellite cells contribute to ectopic fat cell formation in skeletal muscle. Nat Cell Biol. 12:143–152.

Uezumi, A., T. Ito, D. Morikawa, N. Shimizu, T. Yoneda, M. Segawa, M. Yamaguchi, R. Ogawa, M.M. Matev, Y. Miyagoe-Suzuki, S. Takeda, K. Tsujikawa, K. Tsuchida, H. Yamamoto, and S. Fukada. 2011. Fibrosis and adipogenesis originate from a common mesenchymal progenitor in skeletal muscle. Journal of Cell Science. 124:3654–3664.

Vallecillo-García, P., M. Orgeur, S. Vom Hofe-Schneider, J. Stumm, V. Kappert, D.M. Ibrahim, S.T. Borno, S. Hayashi, F. Relaix, K. Hildebrandt, G. Sengle, M. Koch, B. Timmermann, G. Marazzi, D.A. Sassoon, D. Duprez, and S. Stricker. 2017. Odd skipped-related 1 identifies a population of embryonic fibro-adipogenic progenitors regulating myogenesis during limb development. Nat Commun. 8:1218.

van de Wetering, M., Oosterwegel, M., Dooijes, D., and Clevers, H.C., 1991. Identification and cloning of TCF-1, a T cell-specific transcription factor containing a sequence-specific HMG box. EMBO J., 10, 123–132

van de Wetering, M., Sancho, E., Verweij, C., de Lau, W., Oving, I., Hurlstone, A., Van der Horn, K., Batlle, E., Coudreuse, D., Haramis, A-P., Tjon-Pon-Fong, M., Moerer, P., Van den Born, M., Soete, G., Pals, S., Eilers, M., Medema, R., Clevers, H., 2002. The beta-catenin/TCF4 complex imposes a crypt progenitor phenotype on colorectal cancer cells. Cell, 111, 241–250

Vallée, A., Lecarpentier, Y., Guillevin, R., & Vallée, J.-N. (2017). Interactions between TGF-β, canonical WNT/β-catenin pathway and PPARγ in radiation-induced fibrosis. Oncotarget; Vol 8, No 52. Retrieved from http://legacy.oncotarget.com/index.php?journal=oncotarget&amp

Vidal, B., Serrano, A. L., Tjwa, M., Suelves, M., Ardite, E., De Mori, R., … Muñoz-Cánoves, P. (2008). Fibrinogen drives dystrophic muscle fibrosis via a TGFβ/alternative macrophage activation pathway. Genes and Development. https://doi.org/10.1101/gad.465908

Wei, J. et al. Canonical Wnt signaling induces skin fibrosis and subcutaneous lipoatrophy: a novel mouse model for scleroderma? Arthritis Rheum. 63, 1707–1717 (2011).

Weise, A., Bruser, K., Elfert, S., Wallmen, B., Wittel, Y., Wohrle, S. and Hecht, A. (2010) Alternative splicing of Tcf7l2 transcripts generates protein variants with differential promoter-binding and transcriptional activation properties at Wnt/beta-catenin targets. Nucleic Acids Res., 38, 1964–1981.

Wosczyna, M.N., A.A. Biswas, C.A. Cogswell, and D.J. Goldhamer. 2012. Multipotent progenitors resident in the skeletal muscle interstitium exhibit robust BMP-dependent osteogenic activity and mediate heterotopic ossification. J Bone Miner Res. 27:1004–1017.

Wosczyna, M.N., and T.A. Rando. 2018. A Muscle Stem Cell Support Group: Coordinated Cellular Responses in Muscle Regeneration. Dev Cell. 46:135–143.

Wosczyna, M. N., Konishi, C. T., Perez Carbajal, E. E., Wang, T. T.,Walsh, R. A., Gan, Q.,Wagner, M.W. and Rando, T. A. (2019). Mesenchymal stromal cells are required for regeneration and homeostatic maintenance of skeletal muscle. Cell Rep. 27, 2029-2035.e5. doi: 10.1016/j.celrep.2019.04.074

Wynn, T. A., & Ramalingam, T. R. (2012). Mechanisms of fibrosis: Therapeutic translation for fibrotic disease. Nature Medicine. https://doi.org/10.1038/nm.2807

Xiang, F. L., Fang, M., & Yutzey, K. E. (2017). Loss of β-catenin in resident cardiac fibroblasts attenuates fibrosis induced by pressure overload in mice. Nature Communications. https://doi.org/10.1038/s41467-017-00840-w

Yamada, M., Ohnishi, J., Ohkawara, B., Iemura, S., Satoh, K., Hyodo-Miura, J., … Shibuya, H. (2006). NARF, an Nemo-like Kinase (NLK)-associated Ring Finger Protein Regulates the Ubiquitylation and Degradation of T Cell Factor/Lymphoid Enhancer Factor (TCF/LEF). Journal of Biological Chemistry, 281(30), 20749–20760. https://doi.org/10.1074/jbc.M602089200

Yamamoto, H., Ihara, M., Matsuura, Y., & Kikuchi, A. (2003). Sumoylation is involved in β-catenin-dependent activation of Tcf-4. EMBO Journal. https://doi.org/10.1093/emboj/cdg204

Yuan, T., Yan, F., Ying, M., Cao, J., He, Q., Zhu, H., & Yang, B. (2018). Inhibition of Ubiquitin-Specific Proteases as a Novel Anticancer Therapeutic Strategy. Frontiers in Pharmacology. https://doi.org/10.3389/fphar.2018.01080. Retrieved from https://www.frontiersin.org/article/10.3389/fphar.2018.01080

