## Supplemental Table 1 for "TGF-β-driven downregulation of the Wnt/β-Catenin transcription factor TCF7L2/TCF4 in PDGFRα^+^ fibroblasts"

Table S1: Primers used in RT-qPCR

| Gene | Forward primer (5'-3') | Reverse primer (5'-3') |
|---|---|---|
| *Tcf7* | GCCAGAAGCAAGGAGTTCAC | ACTGGGCCAGCTCACAGTAT |
| *Lef1* | CGCTAAAGGAGAGTGCAGCTA | GCTGTCTCTCTTTCCGTGCT |
| *Tcf7l1 (Tcf3)* | TGGTCAACGAATCGGAGAAT | TCACTTCGGCGAAATAGTCG |
| *Tcf7l2 (Tcf4)* | GAGATGAGAGCGAAGGTGGT | CGGCTGCTTGTCTCTTTTTC |
| *Axin2* | ACTGACCGACGATTCCATGT | CTGCGATGCATCTCTCTCTG |
| *Sox9* | CTCCGGCATGAGTGAGGT | TCGCTTCAGATCAACTTTGC |
| *Ccnd1* | CCCAACAACTTCCTCTCCTG | TCCAGAAGGGCTTCAATCTG |
| *Ctnnb1* | AAGGCTTTTCCCAGTCCTTC | CCCTCATCTAGCGTCTCAGG |
| *Nfatc1* | AACGCCCTGACCACCGATAGCACT | CCCGGCTGCCTTCCGTCTCATA |
| *18S* | TGACGGAAGGGCACCACCAG | CACCACCACCCACGGAATCG |
