## Supplemental Figure 1 for "TGF-β-driven downregulation of the Wnt/β-Catenin transcription factor TCF7L2/TCF4 in PDGFRα^+^ fibroblasts"

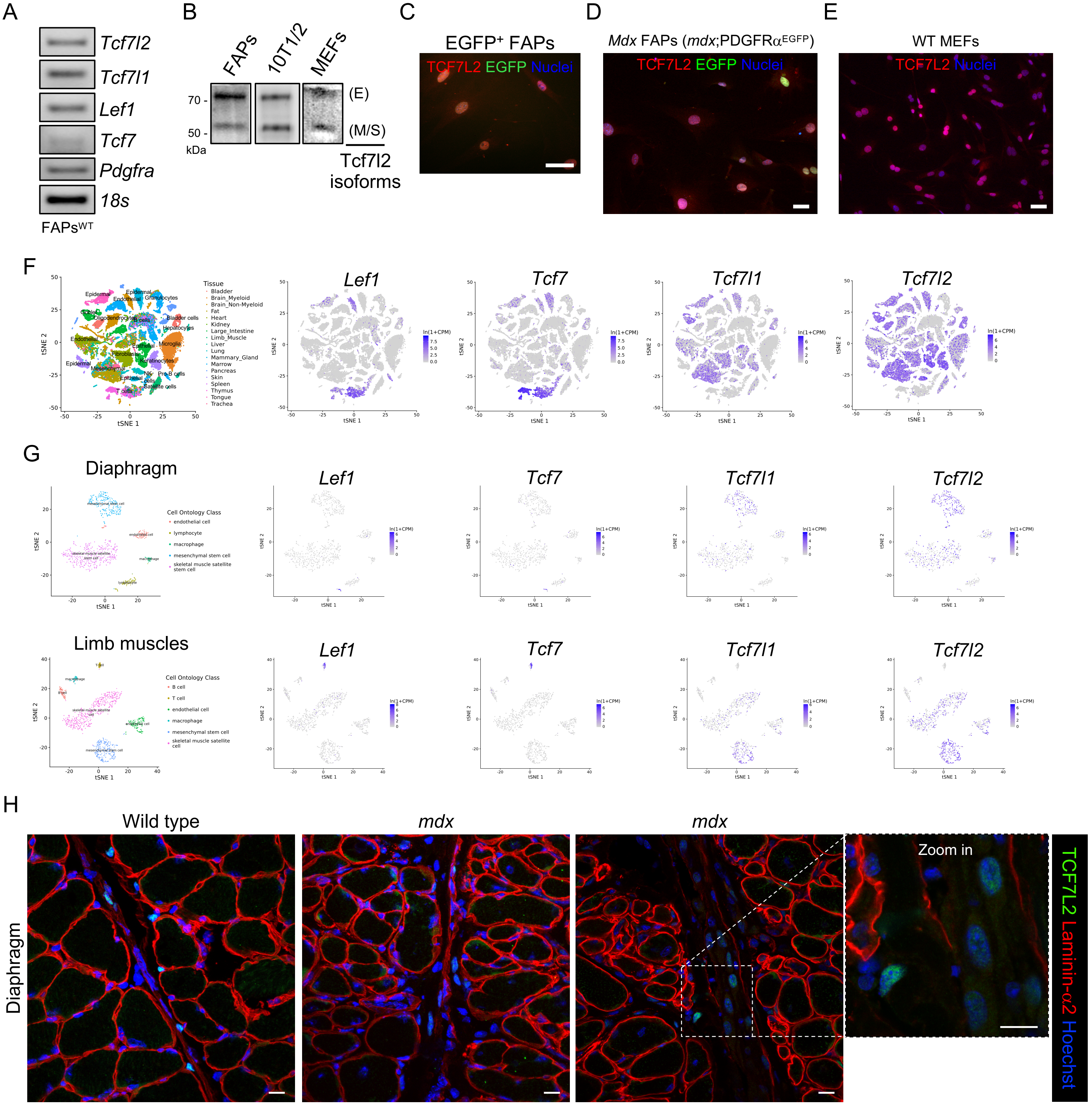


**Fig. S1. Dynamic gene expression of Tcf/Lef in PDGFRα⁺ fibro-adipogenic progenitors and MSCs.**

(A) Final gene expression product of Tcf/Lef was analyzed by RT-qPCR in a 2% agarose gel following 35 PCR cycles. The expression of *Pdgfra* was used as a positive control. (B) Representative western blot analysis showing TCF7L2 protein levels in muscle PDGFRa-EGFP$^+$ FAPs, C3H 10T1/2 MSCs, and MEFs. (C-E) Immunofluorescence of TCF7L2 in FACS-isolated skeletal muscle PDGFRa-EGFP$^+$ FAPs (C), PDGFRa-EGFP$^+$ *mdx* FAPs (D), and cultured MEFs (E). Scale bar: 50 μm. TCF7L2 immunofluorescence (*red*) in MEFs. (F) A t-SNE plot of all cells collected by the microfluidic-droplet method, colored by the predominant cell type that composes each cluster. Cells were colored by cell type for diaphragm and limb muscles and visualized with t-SNE. t-SNE visualization of *Tcf/Lef* genes (from *grey*, low expression, to *blue*, high expression). (H) Z-stack confocal images showing the localization of TCF7L2$^+$ cells in diaphragm muscle sections of adult wild-type and from the dystrophic *mdx* mice. Scale bars: 10 μm.
