## Supplemental Figure 2 for "TGF-β-driven downregulation of the Wnt/β-Catenin transcription factor TCF7L2/TCF4 in PDGFRα^+^ fibroblasts"

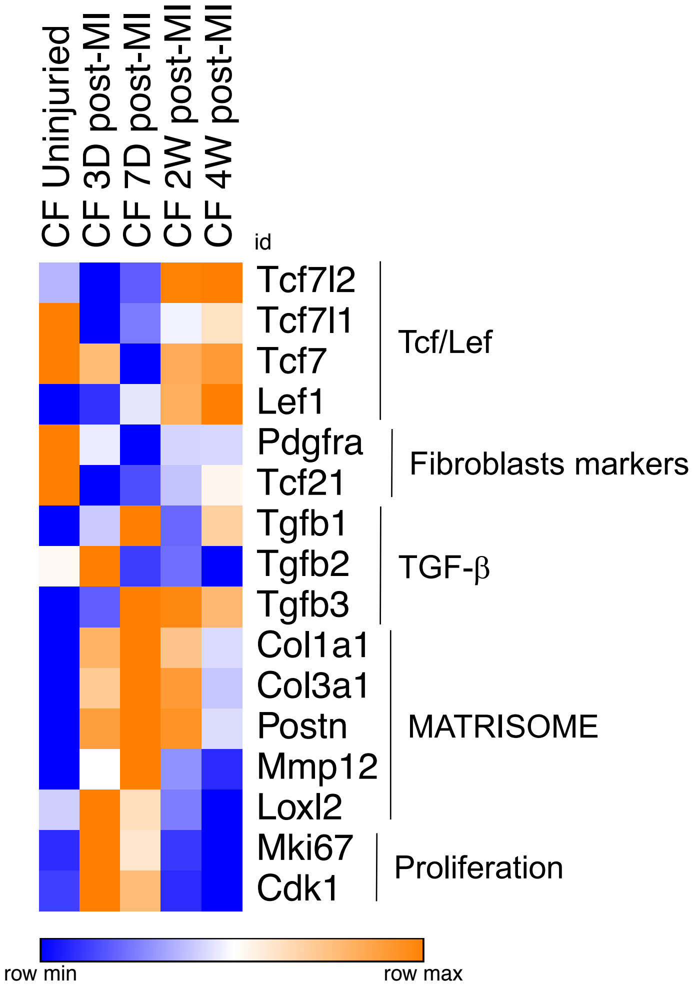


**Fig. S2. Dynamics of Tcf/Lef gene expression in cardiac fibroblasts following myocardial infarction.**

Heat map showing expression changes (RPKMs) of Tcf/Lef genes. Known fibroblasts markers, TGF-β ligands, matrix mediators (e.g. ECM genes), and proliferation-related genes are also shown. Each row is normalized to itself. Each column, per condition, represents the mean of three individual cardiac fibroblast samples (*n*=3).
