## Supplemental Figure 3 for "TGF-β-driven downregulation of the Wnt/β-Catenin transcription factor TCF7L2/TCF4 in PDGFRα^+^ fibroblasts"

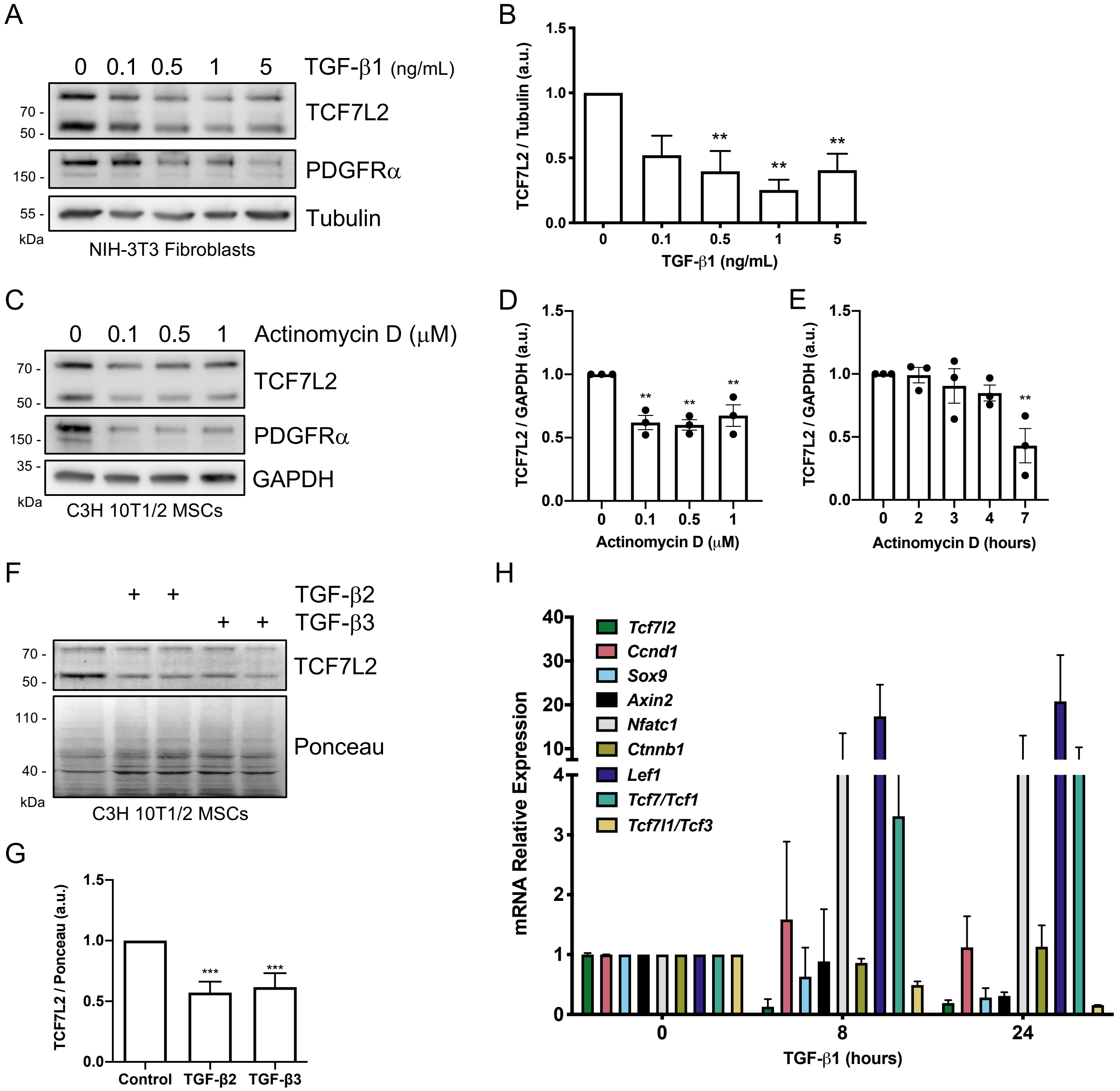


**Fig. S3. Extracellular TGF-β ligands impair TCF7L2-mediated Wnt gene expression.**

(A) Representative western blot analysis showing TCF7L2 and PDGFRα expression levels in NIH-3T3 fibroblasts after treatment with different concentrations of TGF-β1 for 24 h. GAPDH was used as the loading control. (B) Quantification of TCF7L2 protein expression. $**P<0.005$ by one-way ANOVA with Dunnett's post-test; $n=4$. (C) Representative western blot analysis showing TCF7L2 and PDGFRα levels in MSCs after treatment with different concentrations of actinomycin D for 7 h. GAPDH was used as the loading control. (D,E) Quantification of TCF7L2 protein expression. $**P<0.005$ by one-way ANOVA with Dunnett's post-test; $n=3$. (F) Representative western blot from three independent experiments, showing TCF7L2 protein levels after stimulation with TGF-β2 and TGF-β3 for 24 h (5 ng/ml) in MSCs. Ponceau was used as the loading control. (G) Quantification of TCF7L2 protein expression $***P<0.001$; One-Way ANOVA with Dunnett post-test; $n=3$. (H) *Tcf4* (*Tcf7l2*), *Ccnd1* (*CyclinD1*), *Sox9*, *Axin2*, *Nfatc1*, *Ctnnb1* (*β-catenin*), *Lef1*, *Tcf7* (*Tcf1*), and *Tcf7l1* (*Tcf3*) mRNA expression levels were analyzed by quantitative PCR in TGF-β1-treated C3H/10T1/2 MSCs at different time points (0, 8, 24 h). $n=3$.
