## Supplemental Figure 4 for "TGF-β-driven downregulation of the Wnt/β-Catenin transcription factor TCF7L2/TCF4 in PDGFRα^+^ fibroblasts"

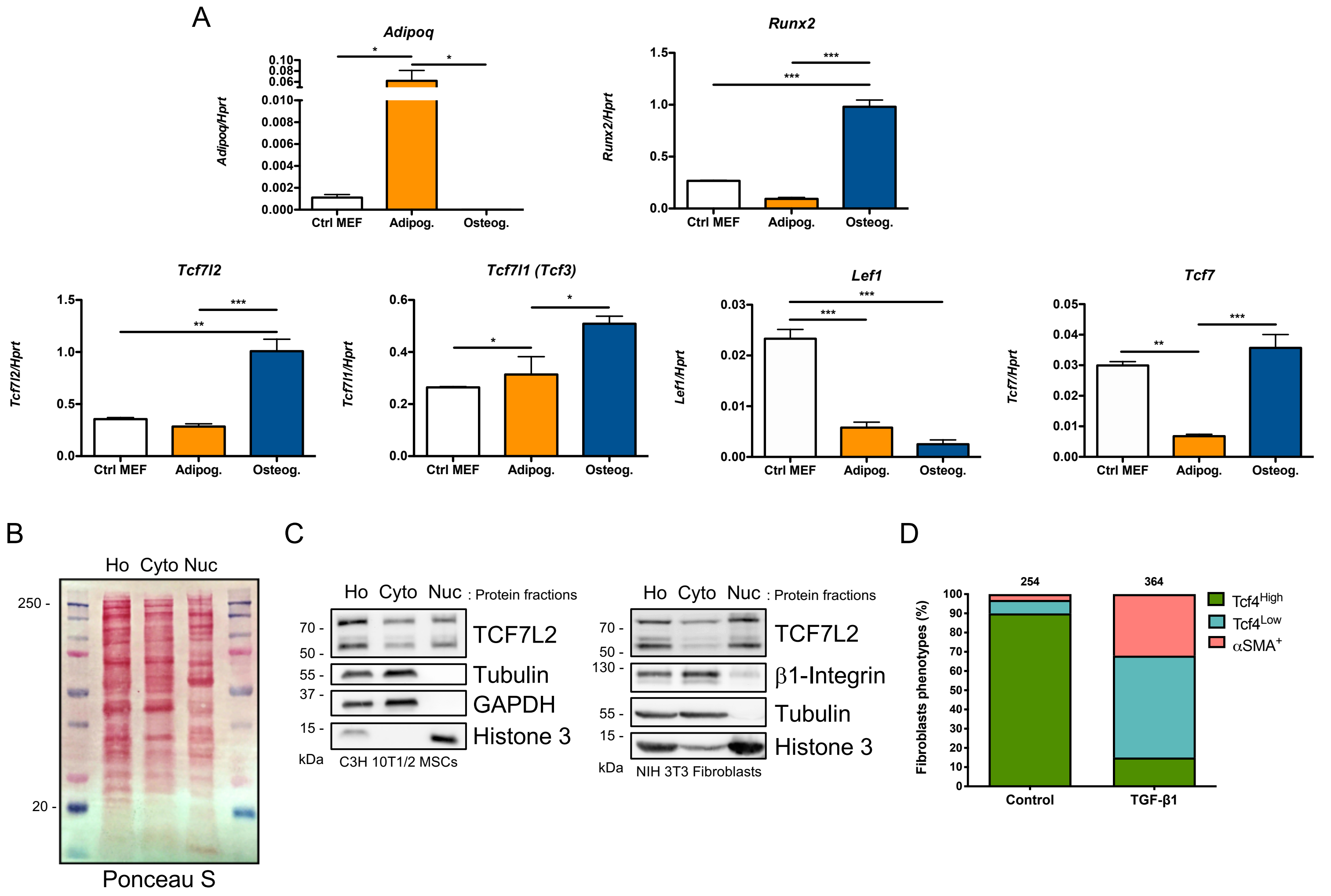


**Fig. S4. Tcf/Lef gene expression varies in adipocytes and osteocytes.**

(A) Adipoq, Runx2, Tcf7l2, Tcf7l1, Lef1, and Tcf7 mRNA expression levels were analyzed by digital droplet RT-qPCR in MEFs control (Ctrl MEF), MEFs-derived adipocytes (Adipog.: adipogenic cell medium), and MEFs-derived osteocytes (Osteog.: osteogenic cell medium). (B) Representative ponceau red staining of Ho: Whole cell lysate; Cyto: Cytoplasmic lysate; Nuc: Nuclei lysate. (C) Representative western blot analysis showing TCF7L2, Tubulin, GAPDH, Histone 3, and β1-Integrin protein levels in proliferating C3H/10T1/2 MSCs and NIH-3T3 fibroblasts. (D) Quantification of TCF7L2 fluorescence intensity in control- and TGF-β1-treated fibroblasts. $TCF7L2^{Hi}$ ($Tcf4^{Hi}$), $TCFL2^{low}$ ($Tcf4^{low}$), and αSMA+-phenotypes were quantified in control and TGF-β1-stimulated C3H/10T1/2 MPCs at 36 h. The numbers above each graph show the total quantified number of cells. n=3.
