## Supplemental Figure 5 for "TGF-β-driven downregulation of the Wnt/β-Catenin transcription factor TCF7L2/TCF4 in PDGFRα^+^ fibroblasts"

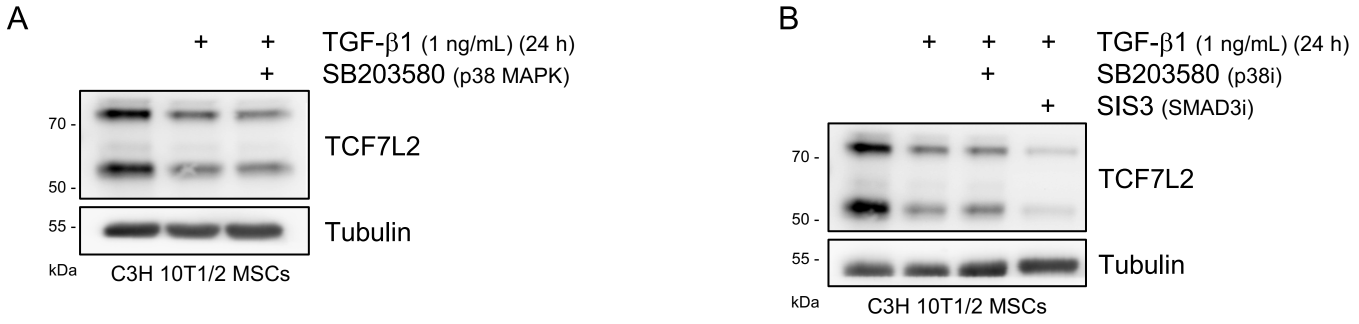


**Fig. S5. Pharmacological Smad3 inhibition with SIS3 pronounces TGF-β-mediated downregulation of TCF7L2.**

(A) Representative western blot analysis showing TCF7L2 expression levels in C3H/10T1/2 cells after TGF-β1 treatment (1 ng/ml) for 24 h. SB203580 (p38 MAPK inhibitor) was co-incubated with TGF-β1 for 24 h. Tubulin was used as the loading control. (B) Representative western blot analysis showing TCF7L2 expression levels in C3H 10T1/2 cells after TGF-β1 treatment (1 ng/ml) for 24 h. SB203580 (p38 MAPK inhibitor) or SIS3 (Smad3 inhibitor) were co-incubated with TGF-β1 for 24 h. Tubulin was used as the loading control.
