## Supplemental Figure 6 for "TGF-β-driven downregulation of the Wnt/β-Catenin transcription factor TCF7L2/TCF4 in PDGFRα^+^ fibroblasts"

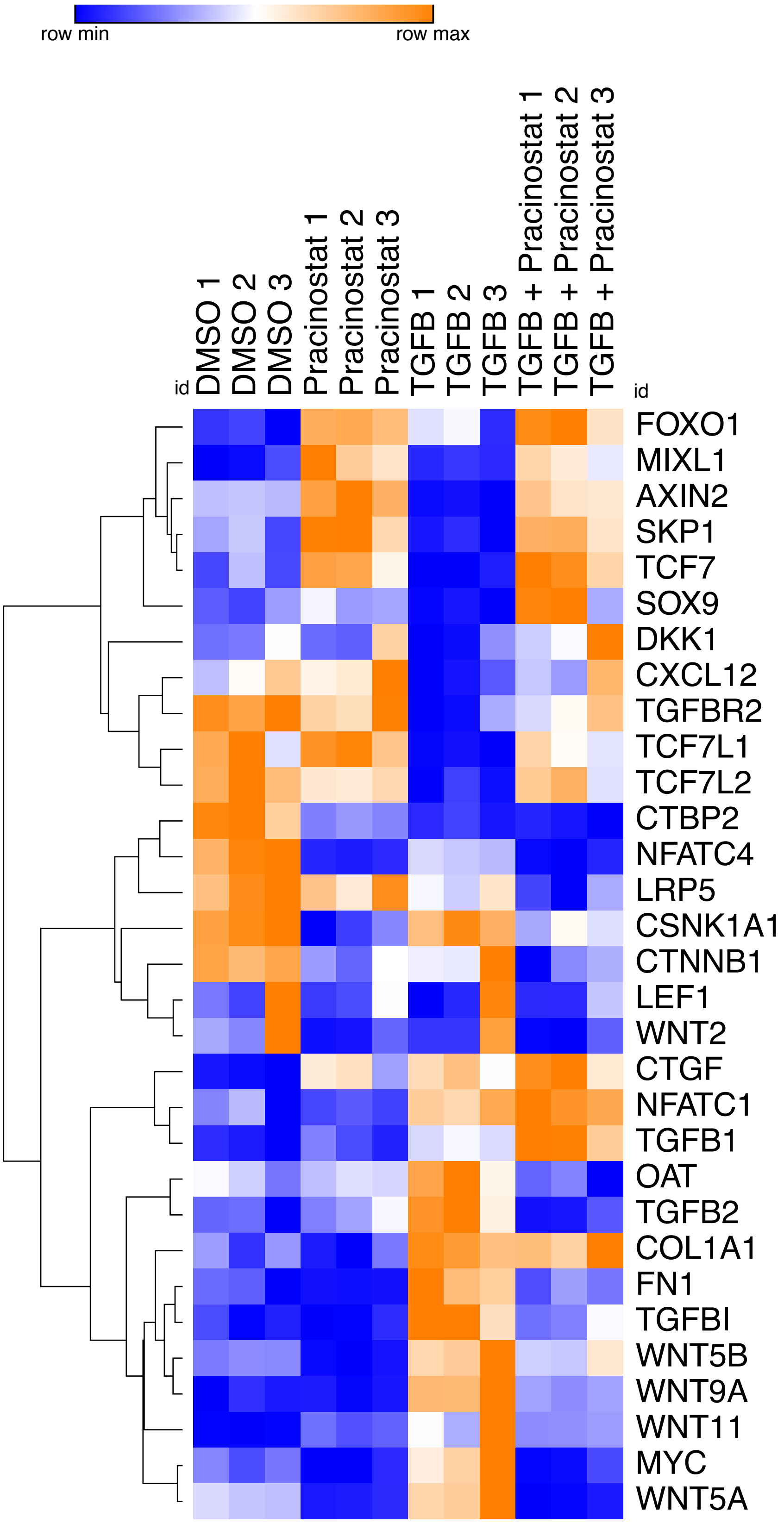


**Fig. S6. TGF-β signaling impairs TCF7L2-mediated target gene expression and Wnt signaling via HDACs.**

Heat map showing expression changes of several validated TCF7L2-downstream genes that are significantly repressed or increased by TGF-β and decreased by the pan HDAC inhibitor pracinostat. Each column, per treatment condition, represents an individual IPF lung fibroblast donor (n=3).
