## Supplemental Figure 7 for "TGF-β-driven downregulation of the Wnt/β-Catenin transcription factor TCF7L2/TCF4 in PDGFRα^+^ fibroblasts"

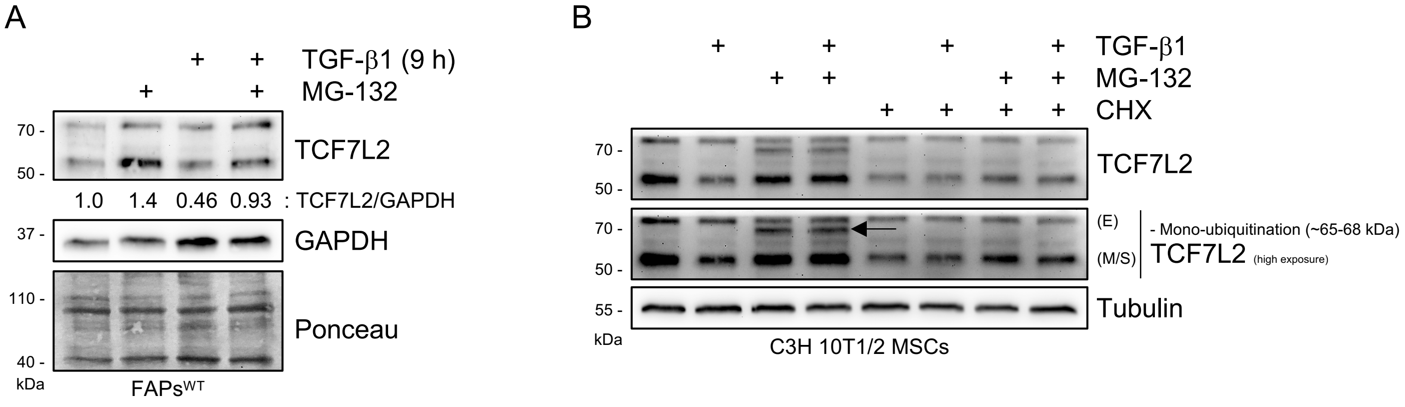


**Fig. S7. Evaluation of the participation of the ubiquitin-proteasome system via MG132 inhibitor on the regulation of TCF7L2 protein expression.**

(A) Representative western blot analysis showing TCF7L2 expression levels in C3H 10T1/2 cells after TGF-β1 treatment (5 ng/ml) for 9 h. MG132 (26S subunit proteasome inhibitor) was incubated alone or co-incubated with TGF-β1 for 9 h. GAPDH and ponceau red were used as the loading control. (B) Representative western blot analysis showing TCF7L2 expression levels in C3H 10T1/2 cells after TGF-β1 treatment (5 ng/ml) for 9 h. MG132 (26S subunit proteasome inhibitor) or cycloheximide (protein translation inhibitor) were incubated alone or co-incubated with TGF-β1 for 9 h. Tubulin was used as the loading control.
