## Supplemental Figure 8 for "TGF-β-driven downregulation of the Wnt/β-Catenin transcription factor TCF7L2/TCF4 in PDGFRα^+^ fibroblasts"

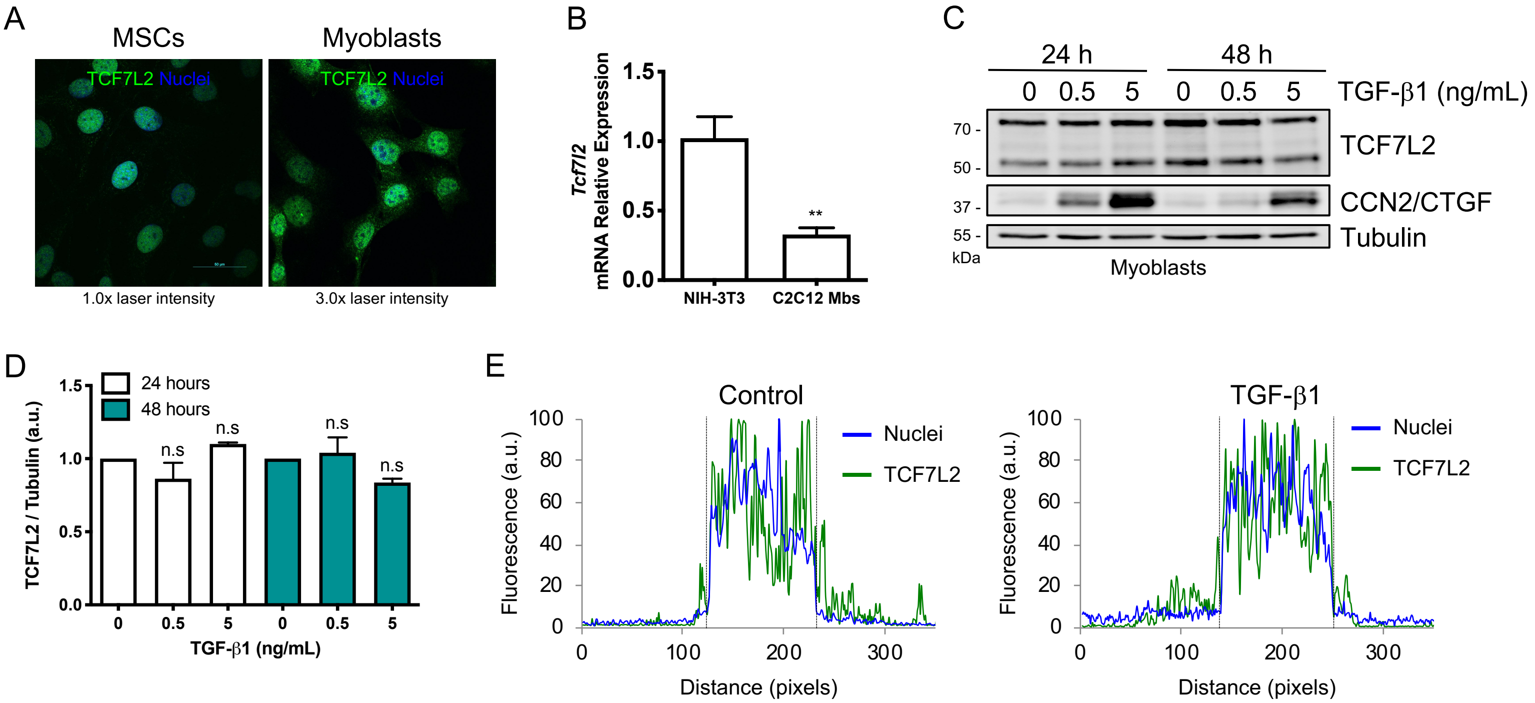


**Fig. S8. The expression of TCF7L2 is not affected by TGF-β signaling in C2C12 myoblasts.**

(A) Confocal images showing TCF7L2 localization in C3H/10T1/2 MSCs and C2C12 myoblasts cell types. Nuclei were stained with Hoechst (blue). Laser intensities (low vs high) were manually adjusted to show similar intensities of TCF7L2 fluorescence in both cell types. (B) *Tcf7l2* mRNA expression levels were analyzed by quantitative PCR in proliferating NIH-3T3 fibroblasts and C2C12 myoblasts. **P<0.005 by two-tailed Student's t-test. n=3. (C) Representative western blot analysis of three independent experiments, showing TCF7L2 and CCN2/CTGF protein levels in TGF-β1-treated C2C12 myoblasts at different concentrations for 24 or 48 h. Tubulin was used as the loading control. (D) Quantification of TCF7L2 protein levels. n.s, not significant by one-way ANOVA with Dunnett's post-test; n=3. (E) Label-distribution graph showing the fluorescence intensity of TCF7L2 and Hoechst along the cell axis. Distance is shown in pixels. Dotted lines show the nucleus-cytoplasm boundary. (a.u.: arbitrary units).
